# Hierarchical encoding of reward, effort and choice across the cortex and basal ganglia during cost-benefit decision making

**DOI:** 10.1101/2023.10.31.563750

**Authors:** Oliver Härmson, Isaac Grennan, Brook Perry, Robert Toth, Colin G. McNamara, Timothy Denison, Hayriye Cagnan, Sanjay G. Manohar, Mark E. Walton, Andrew Sharott

**Affiliations:** Medical Research Council Brain Network Dynamics Unit, Nuffield Department of Clinical Neurosciences, University of Oxford, Oxford, United Kingdom; Department of Experimental Psychology, University of Oxford, Oxford, United Kingdom; Department of Engineering Science, University of Oxford, Oxford, OX1 3PJ, UK; Nuffield Department of Clinical Neurosciences, University of Oxford, Oxford, United Kingdom; Wellcome Centre for Integrative Neuroimaging, University of Oxford, Oxford, United Kingdom; APC Microbiome Ireland, University College Cork, Cork, Ireland; Department of Bioengineering, Imperial College London, London, United Kingdom

**Author notes:** These authors contributed equally to this work (Oliver Härmson, Isaac Grennan, Mark E. Walton, Andrew Sharott). **To whom correspondence should be addressed:** Oliver Härmson; Andrew Sharott.

## Abstract

Adaptive value-guided decision-making requires weighing up the costs and benefits of pursuing an available opportunity. Though neurons across frontal cortical-basal ganglia circuits have been repeatedly shown to represent decision-related parameters, it is unclear whether and how this information is coordinated. To address this question, we performed large-scale single unit recordings simultaneously across 5 medial/orbital frontal and basal ganglia regions as rats decided whether to pursue varying reward payoffs available at different effort costs. We found that single neurons encoding combinations of the canonical decision variables (reward, effort and choice) were represented within all recorded brain regions. Co-active cell assemblies - ensembles of neurons that repeatedly co-activated within short time windows (<25ms) within and across structures - were able to provide representations of the same decision variables through the synchronisation of individual neurons with different coding properties. Together, these findings demonstrate a hierarchical encoding structure for cost-benefit computations, where individual neurons with diverse encoding properties are coordinated into larger, low-dimensional spaces within and across brain regions that can signal decision parameters on the millisecond timescale.

## Introduction

Deciding whether to invest effort in the pursuit of potential rewards is a challenge that organisms continually face. A wide range of areas across the medial frontal cortical-basal ganglia circuit have been implicated in such cost-benefit decisions ^1–9^. Single-units within individual regions have been shown to code decision-relevant variables such as expected reward value and effort cost ^10–20^. However, we currently lack a mechanistic understanding of how activity across these frontal and basal ganglia regions account for how animals evaluate a beneficial opportunity and regulate whether actions are initiated to pursue that opportunity.

One potential reason is that while work to date has focused on how single neurons in individual regions encode key cost-benefit decision parameters, much less is known about how such coding is *coordinated* across an extended network of brain regions. A common proposition is that single neuron responses reflect and contribute to modular processing across specialised brain areas that arrive at a behavioural output through serial information transmission ^21–23^, akin to that described for sensory pathways. In sensorimotor tasks, information from cortex is transferred reliably and topographically to connected areas of striatum ^24^. However, this need not be the case for cost-benefit calculations, where information presented over different timescales needs to be integrated and then utilised to regulate subsequent choice.

One mechanism to achieve such distributed processing is the formation and expression of co-firing assemblies, whereby groups of neurons repeatedly fire action potentials within short-latency (10-30ms) of each other ^25,26^. These assemblies have been postulated to play a vital role in population-level representations, and co-firing assembly patterns in the hippocampus have been repeatedly demonstrated to encode spatial location ^27–29^. In addition, coding by short-timescale coincident spiking can effectively be read out by downstream neural populations on timescales that are optimal for inducing plasticity ^25,30^. Despite the potential of co-firing assemblies to co-ordinate neural activity underlying cost-benefit decision-making, it is not clear whether they are formed in cortico-basal ganglia circuits and, if so, whether they can encode relevant variables. Determining this requires simultaneous large-scale recording of neurons across the network during cost-benefit decision making, which hitherto has rarely been rarely conducted.

To address these questions, we performed simultaneous recordings of single neurons across interconnected frontal and basal ganglia regions – anterior cingulate cortex (ACC), medial/ventral orbitofrontal cortex (MO/VO), dorsal medial striatum (DMS), ventral pallidum (VP) and subthalamic nucleus area (STA) – while rats evaluated whether to accept or reject cost-benefit offers. Individual neurons across the entire network encoded combinations of canonical decision variables (reward, effort and choice), with a particularly pronounced phasic and multiplexed signal in the DMS and STA. Importantly, co-firing assemblies were also detected within and across all cortico-basal ganglia sites encoding the three main decision parameters independently from the individual firing rates of the member neurons. Remarkably, these often included individual neurons whose firing rates signalled a parameter *different* to the tuning of the overall assembly, or neurons with no discernible rate-coding of the canonical decision variables at all. Co-firing assemblies thus coordinate individual neurons with wide-ranging coding properties and give rise to emergent representations that can mediate cost-benefit computations at the network level.

## Results

### Choice on a novel rat accept/reject cost-benefit choice paradigm is shaped by anticipated costs and benefits

To elicit a trade-off between rewards and associated effort costs, we designed a novel cost-benefit task, inspired by foraging scenarios. Rats chose whether to pursue or forgo a reward available at the other end of a linear corridor based on its size and the set level of effort (Fig. 1a, online methods). Trials were initiated by poking and holding their noses in an initiation port (’pre-cue’) while an auditory cue was presented (’offer’), which signalled the size of the available reward on that trial (4 levels: nil, small, medium, large; Fig. 1b). Once the port light illuminated (’Go cue’), rats could then leave the port (’action’) and decide to either (i) pursue the offered reward by running to the opposite end of the corridor (’accept’) or (ii) forgo the offered reward by refraining from entering the corridor (’reject’). In both cases, trials were followed by an inter trial interval (ITI), after which a new trial could be initiated. The effort cost of pursuing the reward was manipulated by placing different numbers of barriers in the corridor, fixed across blocks of ∼20 trials (3 levels: low, moderate, high; Fig 1c).

**Figure 1.**
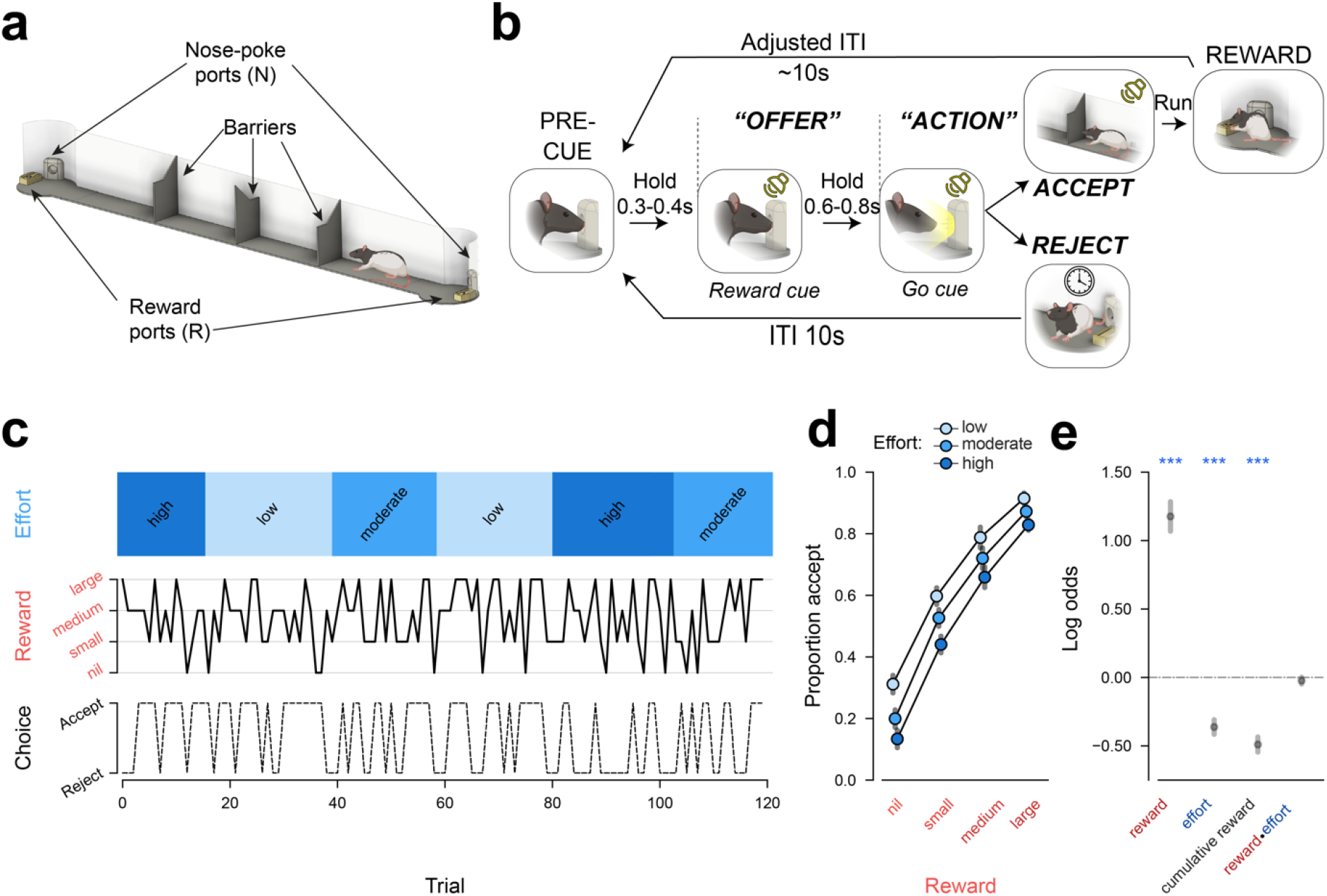
Choice on a novel rat accept/reject choice paradigm is shaped by the costs and benefits. **a**, Illustration of the experimental apparatus which consisted of a 130cm corridor with a variable number of barriers with circular arenas at either end, each of which contained a nose-poke port and a reward port. **b,** Rats chose between reward, signalled by an auditory cue during the “offer” window while the rat was in the nose-poke port, available at the far end of a linear corridor (’accept’) versus opting out (’reject’). To ensure average reward rate was minimally affected by rats’ accept/reject choices, ITIs were adjusted on ‘accept’ trials by subtracting the cohort median run time (online methods). Trials could always be initiated from the same arena where the last trial had finished (same arena on rejected trials; opposite side after an accepted trial). **c,** Reward/effort variation and choice in an example session. Reward magnitude (4 levels, middle panel) is signalled trial-by-trial by an auditory cue, whereas effort (3 levels, upper panel), is expressed in blocks as the number of barriers in the corridor. Choice is depicted as a dotted line on the bottom panel. **d**, Rates of accepting offers as a function of reward (x-axis) and effort (n=12). Coloured dots indicate mean ± SEM, adjusted for the within-subject variance across all levels of effort. **e,** weightings of task variables on behavioural choice from a binomial mixed effects model, depicted as mean ± SEM. Effect of reward x effort: β=-0.01 ± 0.02, p = .69. *** p < .001.

Rats’ choices were sensitive to both the benefits and costs of acting: animals were more likely to pursue reward when it was larger and when the effort cost was lower (n=12, non-implanted animals, Fig. 1c-e). This was reflected as a robust positive influence of reward (β=1.15 ± 0.10, p<.001, binomial mixed effects’ model) and a negative influence of effort (β=-0.33 ± 0.02, p<.001) over choice (Fig. 1e, online methods). Despite the lack of an explicit cue to signal the effort level, it nonetheless had a negative influence on decisions throughout trial blocks (Fig. S1a). Rats also progressively accepted fewer offers over the course of a session as they gained more rewards (β=-0.47 ± 0.02, p<.001), suggesting sensitivity to outcome value (Fig. 1e). This was corroborated by a follow-up experiment where rats (n=6) were given free access to lab chow for 1h prior to testing, which resulted in a significant reduction in willingness to accept any offer (Fig. S1b-c), indicating that choices on this task are not reflective of a fixed decision-making policy.

Sensitivity to reward and effort remained significant in a subset (n=4) of rats who were implanted for electrophysiological recordings in an adapted version of this paradigm (Fig. S1d-e, online methods). Reward offers also consistently influenced action initiation latency on successful trials (Fig. S1f) and premature nose-poke exit rates during the offer cue (Fig. S1j) in both sets of animals. Rats’ performance on this paradigm was thus robustly sensitive to the offered reward magnitude, effort cost and outcome value.

### Cost-benefit decision variable representations are asymmetrically distributed across the frontal cortical-basal ganglia network

Having established that choices on our task were concurrently sensitive both reward and effort, we first aimed to identify neural correlates of the underlying computations at the level of single neurons. Given the role of the frontal-basal ganglia network in motivating cost-benefit choice ^4,9,17,31^, we targeted the medial/ventral orbitofrontal cortex (MO/VO), cingulate gyrus 1/2 (anterior cingulate cortex, ACC), dorsal medial striatum (DMS), ventral pallidum (VP) and subthalamic area (STA, encompassing the subthalamic nucleus, zona incerta, substantia nigra pars reticulata and compacta) (Fig 2a-d, Fig. S2a-e) using a bespoke driveable multielectrode implant to record individual neurons from well-trained rats (n=4) performing the cost-benefit task (online methods).

**Figure 2.**
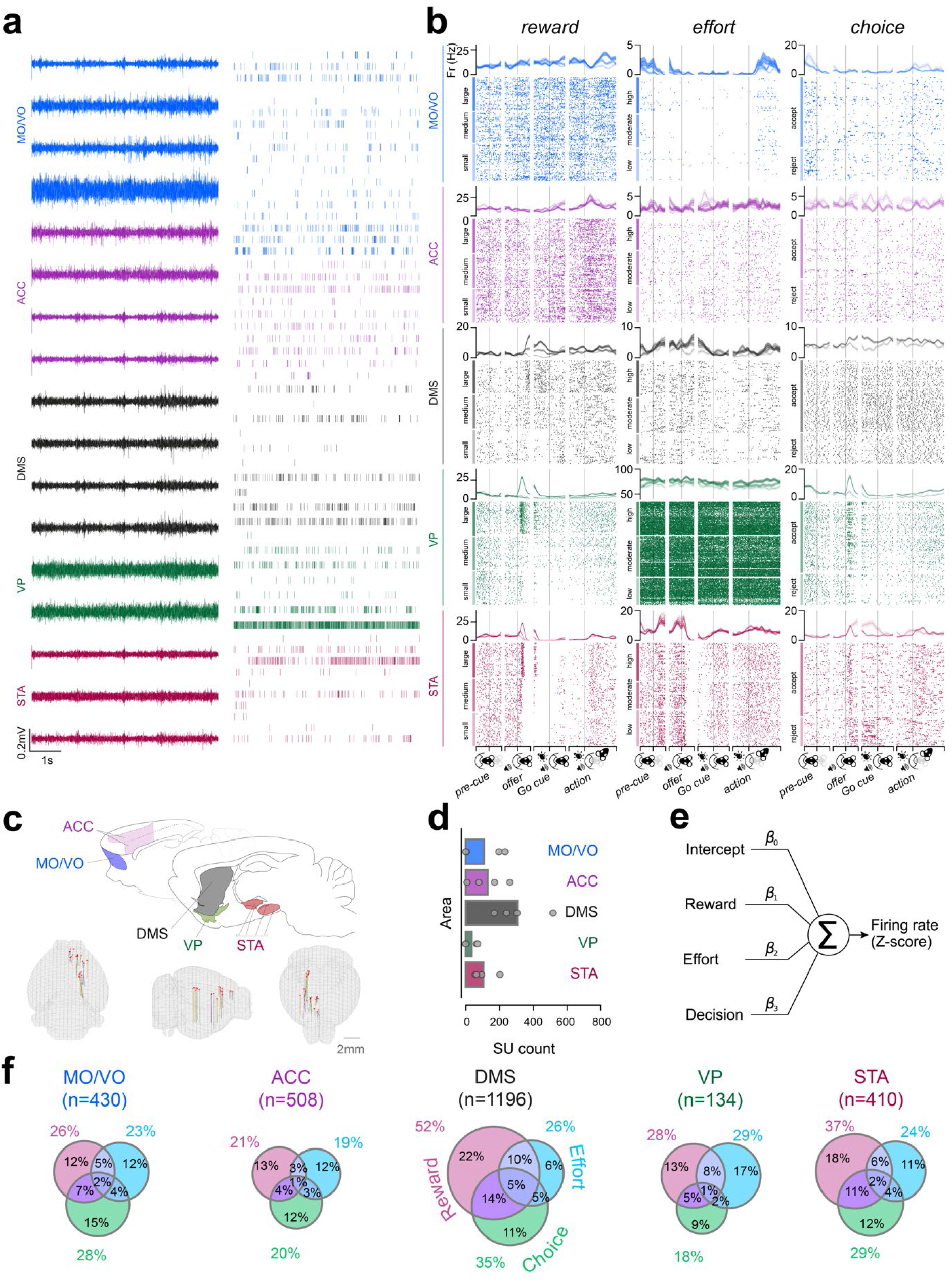
Single-neurons encode combinations of reward, effort, and choice across the frontal cortex and basal ganglia. **a**, High-pass filtered signal (left) and individual neuron spiking times (right, online methods) displayed from an example five-site simultaneous recording from one example subject. **b,** peri-event time histograms (PSTHs, top panels) and raster plots (bottom panels) of 15 individual example neurons (one per panel) that monotonically and significantly varied their firing rate as a function of levels of reward, effort and the rats’ eventual choice (online methods). Data are time-locked to the beginning of the pre-cue, offer, Go cue and action epochs on the x axes, signified by vertical grey bars. PSTH data are displayed as averages across all other task conditions ± SEM (shaded areas). Data are displayed individually for each trial and grouped from top to bottom by levels of a decision variable, indicated on the left by coloured vertical bars. **c,** top: representations of targeted areas on sagittal sections adapted from ^32^; bottom: overview of trode locations on 3D meshes registered to the Allen Common Coordinate Framework 3D space using SharpTrack ^33^, viewed from the back (left), right side (middle) and front (right). **d,** Overview of the average and per-subject single unit (SU) yield in each brain area. Means across subjects are depicted as bars and subject-by-subject cellular yields are marked as grey dots. We isolated on average 108 ± 107 (MO/VO, mean ± standard deviation), 129 ± 96 (ACC), 308 ± 130 (DMS) 34 ± 34 (VP) and 104 ± 58 (STA) single units per animal. **e,** Schematic of the multiple linear regression model used to analyse the contributions of decision variables on individual neuronal firing rates. **f,** Proportions of neurons that were tuned to canonical decision variables across a wide time window in each region (online methods), detected on the basis of significant coefficient of partial determination (CPD) values (p < .050). Pairwise comparisons for reward-coding neuron proportions: DMS vs all other structures: all |z| > 5.13, p < .001, two-sided z-test of proportions, adjusted for multiple comparisons using the BH procedure. STA vs all structures except VP: all |z| > 3.37, p < .001; STA vs VP: |z| = 1.74, p < .050. All other |z| < 1.80, p > .050. Effort-coding neuron proportion comparisons: ACC vs DMS |z| = 3.22, p = .001. ACC vs VP: |z| = 2.69, p = .007; all other |z| < 1.91, p > 050. Pairwise comparisons for choice-coding proportions: DMS vs other regions: all |z| > 2.47, p < .013; STA vs ACC and VP: both |z| > 2.48, p < .012. MO/VO vs ACC and VP: both |z| > 2.31, p < .020. All other |z| < 0.61, p > .050. Pairwise comparisons of the summed percentages of all multiplexing neuron types (intersecting Venn diagram areas) between regions: DMS vs all other regions: all |z| > 3.91, p < .001; STA vs MO/VO and ACC: both |z| > 2.06, p < .020; all other |z| < 1.46, p > .072, one-sided z-test of proportions without BH correction. MO/VO = medial/vental orbitofrontal cortex, ACC = anterior cingulate cortex, DMS = dorsal medial striatum, VP = ventral pallidum, STA = subthalamic area. Note that the for a given decision variable in DMS and STA, the proportion of neurons encoding that variable in additon to others is bigger that the proportion of neurons encoding that variable.

We first examined the propensity of cortical and/or basal ganglia neurons to encode single or multiple decision variables. We modelled each identified neuron’s z-scored firing rate as the weighted sum of offered reward, effort and subsequent accept/reject choice (Fig. 2e, online methods) during epochs containing the *entire decision process* (from the offer cue onset to either corridor entry for accepted trials or the session median corridor entry for rejected trials). Significant proportions of units in each area linearly signalled each canonical decision variable (Fig. 2f, all p < .001, one-sided binomial test against a chance rate of 5% with Benjamini-Hochberg adjustment for multiple tests). Strikingly, reward and choice tuning was more frequent in the DMS and STA compared to other structures, (Fig. 2f, see legend for statistical comparisons). Moreover, the co-representation of decision parameters within single neurons was also more common in DMS and STA compared to either frontal cortical region (Fig. 2f, see legend for statistical comparisons). Together, these data suggest that single neuronal representations are highly distributed, but that there are also significant regional asymmetries, with particular specialisation in the DMS and STA.

### The dynamics of cost-benefit representations in cortex and basal ganglia are underpinned by diversity in temporal and directional properties

Our finding that basal ganglia neurons were more specialised than those in the frontal cortex argues against models of cortico-basal ganglia circuits in which abstract decision variables (subjective reward value and effort cost) are computed in cortical regions and used to regulate action initiation and invigoration via basal ganglia structures^22,34^. Nonetheless, such organisation could be more evident over finer timescales. We thus analysed the coefficients of partial determination (CPD) with finer temporal resolution for each canonical decision variable, neuron, and area (Fig. 3a, online methods). To characterise the temporal properties of the CPD signal, we further analysed (1) CPD-“phasicness” - an index of the dispersion of encoding in each area’s single neuron population over time (online methods) - and (2) latency to maximal CPD.

**Figure 3.**
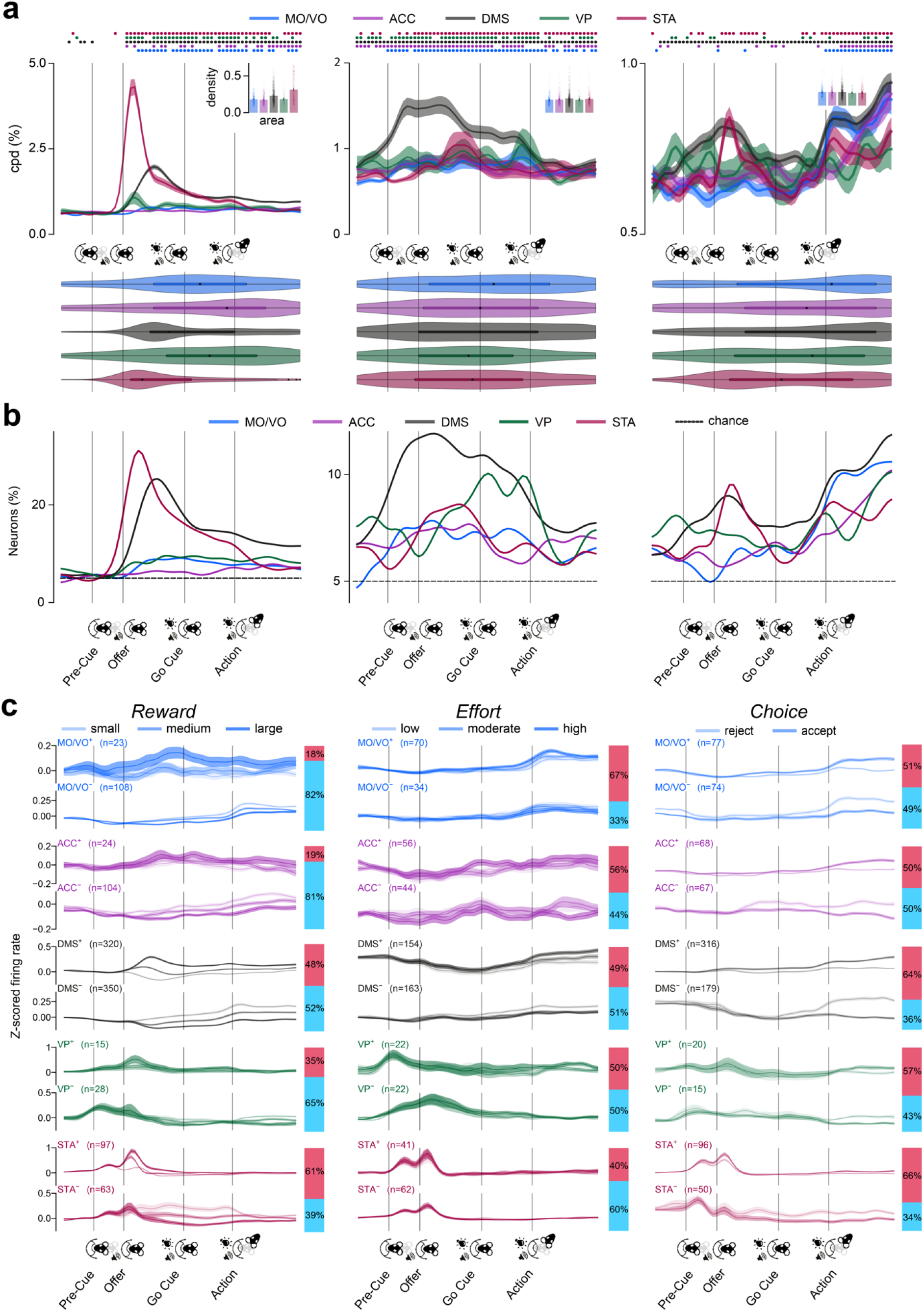
Dynamic and distributed single neuronal representations of reward, effort and choice across the frontal-basal ganglia network. **a**, upper, population average CPD time series, depicted as means (solid lines) ± SEM (shaded areas) across all units, for each decision variable (columns). Colour-coded segments above indicate time points where the population average CPD was significantly higher than expected by chance in each region using a significance threshold of p < .05 (permutation testing, fdr corrected across time points). Insets depict individual unit (coloured dots) and population average (bars) ± SEM phasicness scores for each area. Phasicness comparisons for reward: STA vs all other regions: all |z| > 5.71; p < .001, two-sided Mann-Whitney U test. DMS vs all other regions: all |z| > 3.16 p < .002. All other |z| < 0.58, p > .050. Choice phasicness tests: MO/VO: z = 4.12, p < .001; ACC: z = 3.48, p < .001; DMS: z = 10.18, p < .001; STA and VP: both z < 0.93, p > .05. Lower, latencies to peak CPD, depicted as kernel density estimates (violins) and superimposed boxplots. Instances of peak reward CPDs occurring before offer cue onset were detected at below-chance rate (< 3.72%, p > .05, one-sided binomial test). Reward CPD phasicness tests against bootstrapped null distribution (online methods): MO/VO: z = 1.28, p > .050; ACC: z = 2.27; p = .023; DMS: z = 2.12, p = .027; VP: z = 0.49, p > .050; STA: z = 6.80, p < .001. **b,** Instantaneous recruitment of neurons dedicated to signalling any decision variable with a significance cut-off of p < .05. **c,** Population-averaged z-scored firing rates, grouped by positive (above) or negative (below) tuning valence and plotted on each level of a decision variable (dark lines) ± SEM (shaded areas). Data in each group are averaged across other conditions. Curves in a-c are depicted over a warped trial timeline beginning 0.4s before nose-poke entry up to the median corridor entry time (online methods) and are smoothed with a gaussian filter with a standard deviation of 1.0.

Prospective reward was significantly encoded from the onset of the reward offer cue until the decision point in all areas (Fig. 3a, upper left). Nonetheless, the reward signal in ACC, DMS and STA units was classified as phasic, peaking after presentation of the Go cue, while the VP and MO/VO encoding was more dispersed (ACC, DMS and STA phasicness tests: all |z| > 2.18, p < .029, two-sided Mann-Whitney U test; Fig. 3a). Peak reward CPD and recruitment of neurons was reached earlier in the STA compared to all other areas (Fig. 3a, b left; STA maximum CPD latency vs other areas: |z| > 3.55, p < .001), Whereas the time course of reward coding was clearly linked to the onset of the offer cue, effort signalling was more sustained, being evident in the pre-cue and even pre-trial period in most structures (all CPD phasicness tests, |z| < 1.34, p > .050; Fig. 3a, middle), with similar density (all |z| < 1.08, p > .050) and peak CPD latency (Fig. 3b middle; all |z| < 1.60, p > .050) in all. Choice, on the other hand, elicited phasic firing rate modulation in the MO/VO, ACC, and DMS (Fig. 3a, right; MO/VO, ACC and DMS CPD phasicness tests: |z| > 3.47, p < .001; all other |z| < 0.93, p > .050) with the peak CPD weighted towards the action epoch in all regions (all between-area |z| < 1.72; p .050 for both, Fig. 3b right).

We conjectured that these phasic vs. distributed signals could be underpinned by groups of neurons that differ in their coding valence with respect to the decision parameters^35,36^. As can be seen in Fig. 3c, distributed reward coding in all areas appeared to be driven predominantly by *negatively* coding neurons, while phasic coding in the DMS and STA was mediated mostly by *positively* tuned single units (Fig. 3c, Fig. S3-4). In line with this, the maximum CPD latencies for reward were significantly longer in negative-coding DMS and STA neurons compared to positive ones (comparison of population-median maximum cpd latencies: both |z| 5.31, p < .001, Mann-Whitney U test). Such temporally disperse coding can be underpinned by either (1) persistent temporally disperse activity within single units or (2) tiling of task time by phasic signals of individual neurons (Guo et al. 2017; Runyan et al. 2017). Examination of the phasicness of individual positively vs. negatively reward-coding units revealed that the latter were significantly less phasic and thus more persistently active than the former in the DMS and STA (both |z| > 3.40, p < .001, one-sided Mann-Whitney U test), while phasicness did not differ between these populations in other regions or for other task variables (all other |z| < 1.64, p > .050).

Taken together, these data demonstrate that the temporal dynamics of rate-based encoding varied with respect to brain area, direction and variable. Notably, positively encoding neurons in basal ganglia structures had particularly strong phasic encoding of prospective reward during the offer window, but the peak encoding of the majority of populations was spread across the entire task.

### Neurons across the cortico-basal ganglia network form cell assemblies during cost-benefit decision-making

Such diversity in the single neuronal firing rate read-outs poses a challenge at the output interpretation level. How can levels of reward, effort, and intention to accept/reject these offers be inferred from such spatiotemporally varying inputs? One candidate mechanism of coordination across varying single neuronal signals is the formation of *co-active assemblies*, defined as groups of neurons that synchronise their activity in short time windows. The activity patterns of such assemblies have been proposed to reflect the underlying synaptic connectivity of the constituent neurons and provide a higher-order level of computation ^25^. We thus examined whether cell assemblies could be detected in the cortico-basal ganglia network during cost-benefit decision-making and, if so, whether they could encode decision-relevant variables through the coordination of diverse populations.

We first addressed whether such short-latency cofiring was a common feature of these brain areas during our task. Many pairs of units across all anatomical structures were highly cross-correlated within a short interval (+/- 50 ms) around 0s lag (Fig. 4a-b), consistent with a physiologically meaningful readout for downstream neurons ^25^. These short-latency correlations were dependent on the precise temporal relationship between spike trains, as randomly jittering the spike trains between 0 to 250ms reduced the baseline to peak difference in cross correlation by 71% (Fig. 4c). We used principal component analysis (PCA) to determine the extent of spike timing coordination across groups of simultaneously recorded cells. The added variance explained by the first principal component was much larger for the unjittered z-scored binned spike train (Fig. 4d), as was the mean of the variance added by the first 5 principal components (Fig. 4d). This indicates that a significant proportion of neural activity in the frontal cortical-basal ganglia network is coordinated at the fine temporal scale necessary for the formation and expression of cell assemblies ^25,28,30,37^.

**Figure 4.**
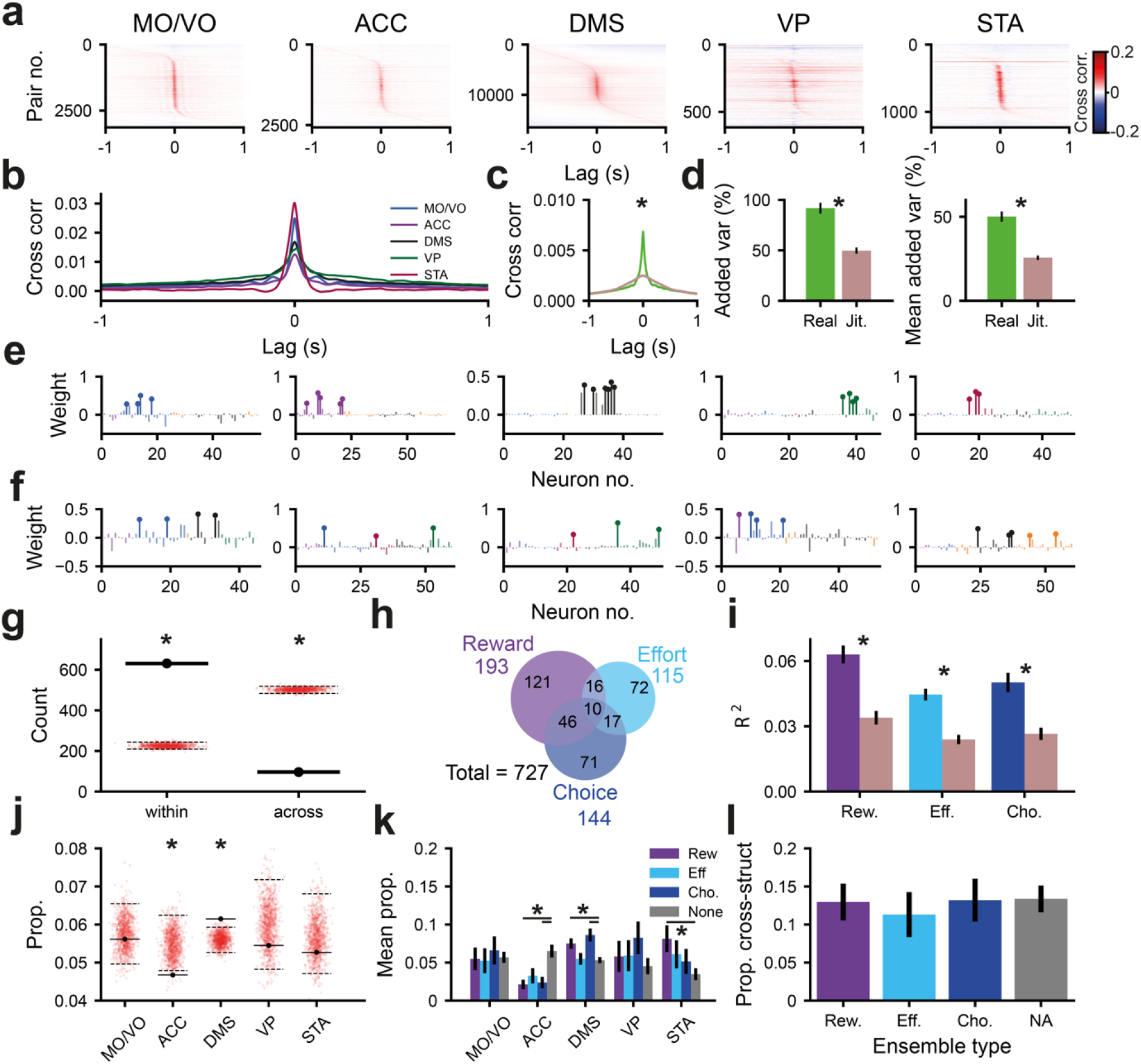
Cofiring assemblies across the frontal-basal ganglia network represent combinations of reward, effort and decision. **a**, heat map showing the cross correlation between pairs of units in the MO, ACC, DMS, VP and STA. **b,** the mean cross correlation between pairs of units in the MO, ACC, DMS, VP and STA. **c,** The mean cross correlation between pairs of units across all structures with (orange) or without (blue) random jittering of spike trains in the range -250ms to 250ms (0ms peak height, p<10^-13^, Wilcoxon signed-rank test). **d,** The mean variance added over that of a single unit for the first eigenvector (left, p<10^-12^, Wilcoxon signed-rank test) and mean variance added of the first 5 eigenvectors (right, p<10^-12^, Wilcoxon signed-rank test) with or without random jittering of spike trains in the range -250ms to 250ms **e,** lollipop plot showing example assembly patterns with members solely within the MO/VO, ACC, DMS, VP and STA (left to right). Member neurons are marked with a circle on top of the bar indicating their weight. Color scheme as in **b**. **f,** example assembly patterns that span multiple anatomical regions. Member neurons are marked with a circle on top of the bar indicating their weight. Color scheme as in **b**. **g,** the number of assembly patterns that span multiple anatomical structures (black marker) as compared to the proportion expected due to chance when shuffling the anatomical labels of neurons (red dots). The black dashed lines mark the 2.5^th^ and 97.5th percentile of this shuffled distribution. **h,** Venn diagram showing the number of assemblies which have their coactivity significantly modulated by reward, effort and decision outcome. **i,** the variance in reward (p<10^-19^, Wilcoxon signed-rank test), effort (p<10^-13^, Wilcoxon signed-rank test) or decision outcome (pseudo-R^2^, p<10^-15^, Wilcoxon signed-rank test), explained by assembly pattern expression strength with (brown) or without (blue) random jittering of spike trains in the range of -250ms to 250ms for significantly encoding assemblies **j,** the mean proportion of SUs from different brain regions which were members of each assembly as compared to the mean proportion when anatomical labels were shuffled amongst neurons (red dots). The 2.5^th^and 97.5^th^ percentiles for shuffled distributions were marked with dashed black lines. **k,** the mean proportion of SUs from different brain regions in assemblies separated by their encoding properties. The proportion of neurons from each anatomical structure in encoding assemblies was compared to assemblies which did not encode canonical variables (significance for p < .050 determined by permutation test, Benjamini-Hochberg procedure used to control for multiple comparisons). **l,** the proportion of cross-structural assemblies by assembly encoding type (p>0.05 for reward, effort and decision vs non-coding, z-test). * p < .050.

Next we identified coactivating assemblies of neurons were using PCA followed by independent component analysis (PCA-ICA) on the binned spike trains of neurons from all recorded regions during a time window spanning the start of the offer cue and the end of the action window (online methods; ^38,39^). This approach yielded a total of 727 assemblies. Most had ‘member neurons’ (i.e., neurons with a high weighting in the assembly pattern) restricted to a single brain region (∼87% Fig. 4e, g). The remaining assemblies had member neurons that spanned multiple structures (Fig. 4f, g), including a small proportion (12/727) which were made up of member neurons spanning 3 or more brain regions (e.g., Fig. 4f). The proportion of assemblies that had member units from within a single structure was significantly greater than chance (Fig. 4g).

We then examined whether such assemblies provide a unified read-out of canonical decision variables. To this end, the mean expression of the coactivity pattern was modelled as a weighted linear sum of reward, effort and choice (online methods). Strikingly, nearly half (49%) of all identified assemblies had a significant degree of selectivity for the prospective reward, current effort cost and choice, either individually or conjointly (Fig. 4h). Jittering the spike trains preserved the mean firing rates over the trial and almost all the rate-coding within the window of analysis (Fig. S5), but substantially disrupted coding at the level of the assembly (Fig. 4i). Overall, assemblies thus provided a substrate of encoding that coordinates activity across populations of neurons but is independent of the underlying correlated changes of firing rate at task-relevant timescales.

To understand how each region contributed to assemblies, we examined the likelihood of neurons from any structure forming assemblies regardless of their underlying tuning properties. We found that DMS and ACC neurons were more or less likely, respectively to be members of co-active assemblies (Fig. 4j). We then examined whether neurons from each area were more likely to be members of assemblies coding for one of the canonical decision variables (blue/purple bars, Fig. 4k) and of assemblies that did not code for any of the decision variables (gray bars, Fig. 4k).

Though the membership of task-tuned assemblies were relatively even across regions, DMS and STA neurons were more likely to be members of reward-coding assemblies than those that did not encode any decision variable (Fig. 4k). DMS neurons were additionally more likely to be members of assembles that encoded choice than those that did not encode any variable. In contrast, ACC neurons were *less* likely to be member of assemblies encoding reward and choice that assemblies that did not encode any variable (Fig. 4k). Cross-structural assemblies were not significantly more likely to encode any of the decision variables (Fig. 4l).

Together, this demonstrates that assemblies were formed by neurons in each structure across the entire network, with DMS neurons in particular having an important role in the expression of reward- and choice-related co-activity.

### Assemblies that encode decision variables are comprised of individual neurons with diverse tuning properties

The above analyses demonstrate that both individual neurons *and* assemblies encode and integrate the canonical decision variables. One possibility therefore is that the assembly code is closely correlated to each individual neuron’s tuning selectivity. However, it is also technically feasible for the temporal code within an assembly to be distinct from the rate code exhibited by the members of that assembly.

To address this, we asked whether ensembles that encoded a given decision variable were comprised of neurons that *rate-coded* the same variable. Strikingly, we found examples of assemblies where the rate-coding of individual neurons clearly diverged completely from that of the overall ensemble (Fig. 5a-f). To quantify the intersection of the encoding properties of single neurons (Fig. 2f) and assemblies (Fig. 4h), we defined the proportion of each type of rate-coding neuron that contributed to each type of encoding assembly (Fig. 5g). Although single units that encoded a given variable were more likely to contribute to assemblies that encoded the same variable than any other assembly type (Fig. 5g), at least 20% of units coding a particular decision variable *also* contributed to assemblies encoding either of the other two variables (Fig. 5g). Only a small proportion of the variance in the assembly pattern expression strength dynamics (<18%, for reward, effort and choice) could be explained by changes in the firing rate of member neurons (Fig. 5h), demonstrating that this bias was not simply a consequence of correlated, task-related firing rate across assembly members.

**Figure 5.**
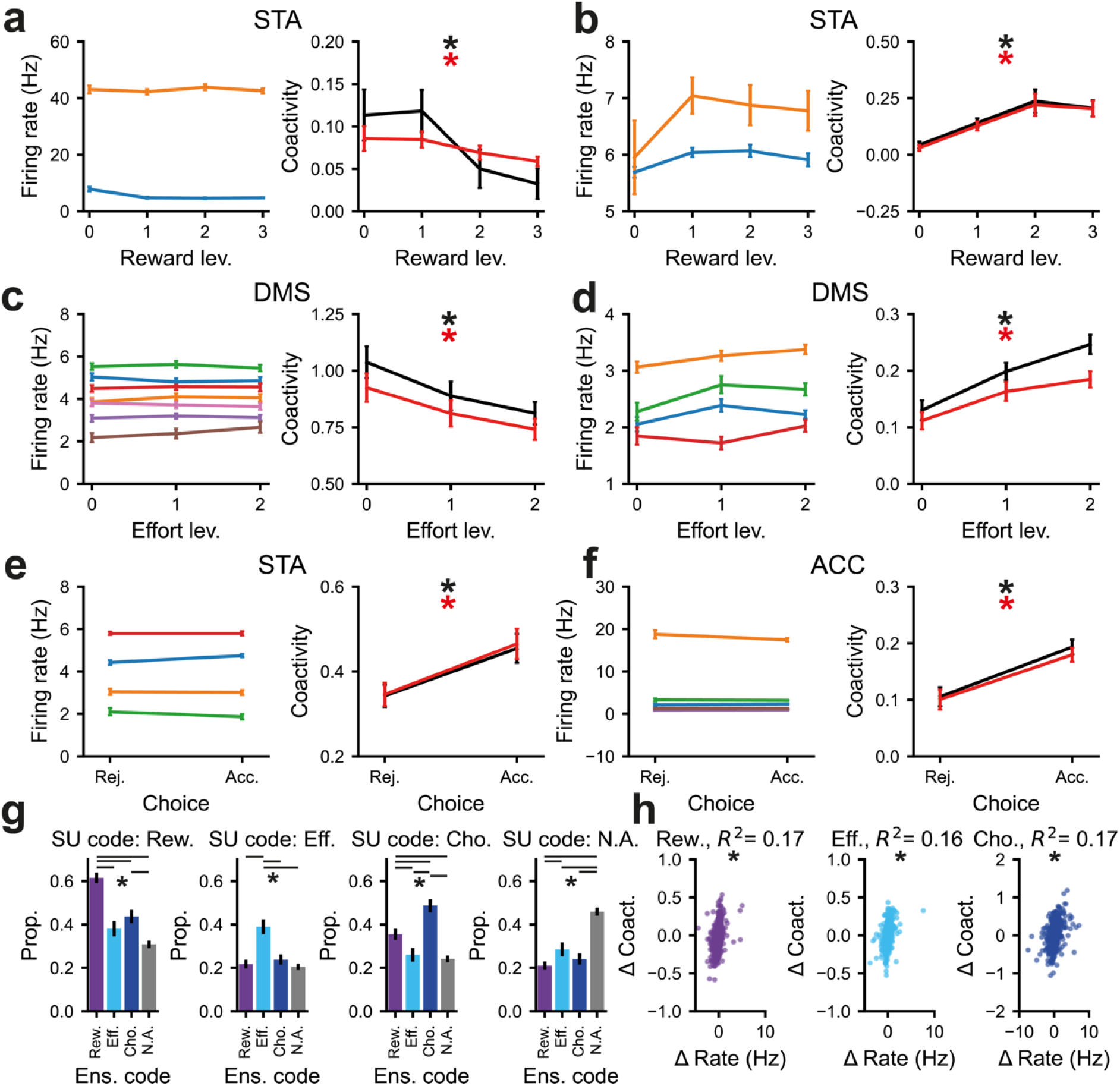
Decision-related encoding by co-firing assemblies can be orthogonal to the rate coding of the individual neurons that comprise them. **a, b**, The firing rate of the member neurons that make up assemblies (left). They show no significant change across reward levels. Assembly coactivity (right) computed using the full assembly pattern weights (black) or restricting weights to only member neurons (red). Both the coactivity for the full assembly pattern and for members only is significantly shaped by the level of offered reward. c**, d,** The firing rate of the member neurons that make up assemblies (left). They show no significant change across effort levels. Assembly coactivity (right) computed using the full assembly pattern weights (black) or restricting weights to only member neurons (red). Both the coactivity for the full assembly pattern and for members is significantly determined by the level of effort. **e** and **f,** The firing rate of the member neurons that make up assemblies (left). They show no significant change with choice outcome. Assembly coactivity (right) computed using the full assembly pattern weights (black) or restricting weights to only member neurons (red). The choice outcome has a significant effect on the coactivity pattern of both the full and members-only assembly pattern. For a-f, * p<.050, significance was determined using linear regressions with firing rate or coactivity as the dependent variable. The CPD of reward, effort and choice outcome was compared to the null distribution from trial shuffled data. **g,** the proportion of member neurons of reward, effort, decision and non-encoding assemblies which rate code reward (left), effort (middle-left), decision outcome (middle-right) or do not rate code for the canonical variables (right). * p <.050, z-test, Benjamini-Hochberg procedure used to control for multiple comparisons. **h,** the change in the firing rate of member neurons across the levels of reward (**i**) and effort (**ii**) and decision (**iii**) outcome only explains a small proportion (<20%) of the change in assembly expression strength. * p < .050 in linear regression.

Taken together, these findings suggest that though a member neuron’s individual tuning often aligns with that of their assembly, assembly pattern expression strength changes were poorly explained by firing rate dynamics. Furthermore, neurons often contributed to assemblies encoding a different parameter to their own. Co-activity is thus separable from rate code and can integrate units with diverse tuning properties.

### Encoding of reward, effort and choice by cortico-basal ganglia assemblies aligns with relevant task events

Our analyses of assemblies so far defined encoding using expression strength across the entire task window. If assemblies provide population-level encoding that has ongoing influence on cost-benefit decisions akin to that proposed for single neurons, their temporal dynamics should align with key task epochs.

To examine whether this was the case, we analysed the time course of encoding (as measured by CPD) for significant *assemblies* with respect to the mean onset of the reward offer, action initiation and nose-poke exit and subsequent accept/reject decision. Significance at a given timepoint was defined in relation to assemblies constructed from jittered spike trains (Fig. S5; online methods). Reward and effort were encoded at above-chance levels during virtually all time-bins from the reward offer until the end of the decision point, whereas choice-encoding assemblies were significant from shortly after the nose-poke exit to the trial end (Fig. 6a, b). As seen for rate-coding of individual neurons, assemblies could encode positively (e.g., maximum expression strength for maximum reward) or negatively (e.g., maximum expression strength for minimum reward). Interestingly, the times course of assemblies that negatively encoded reward was more sustained/less phasic than for positively encoding assemblies (Fig. 6c, d).

**Figure 6:**
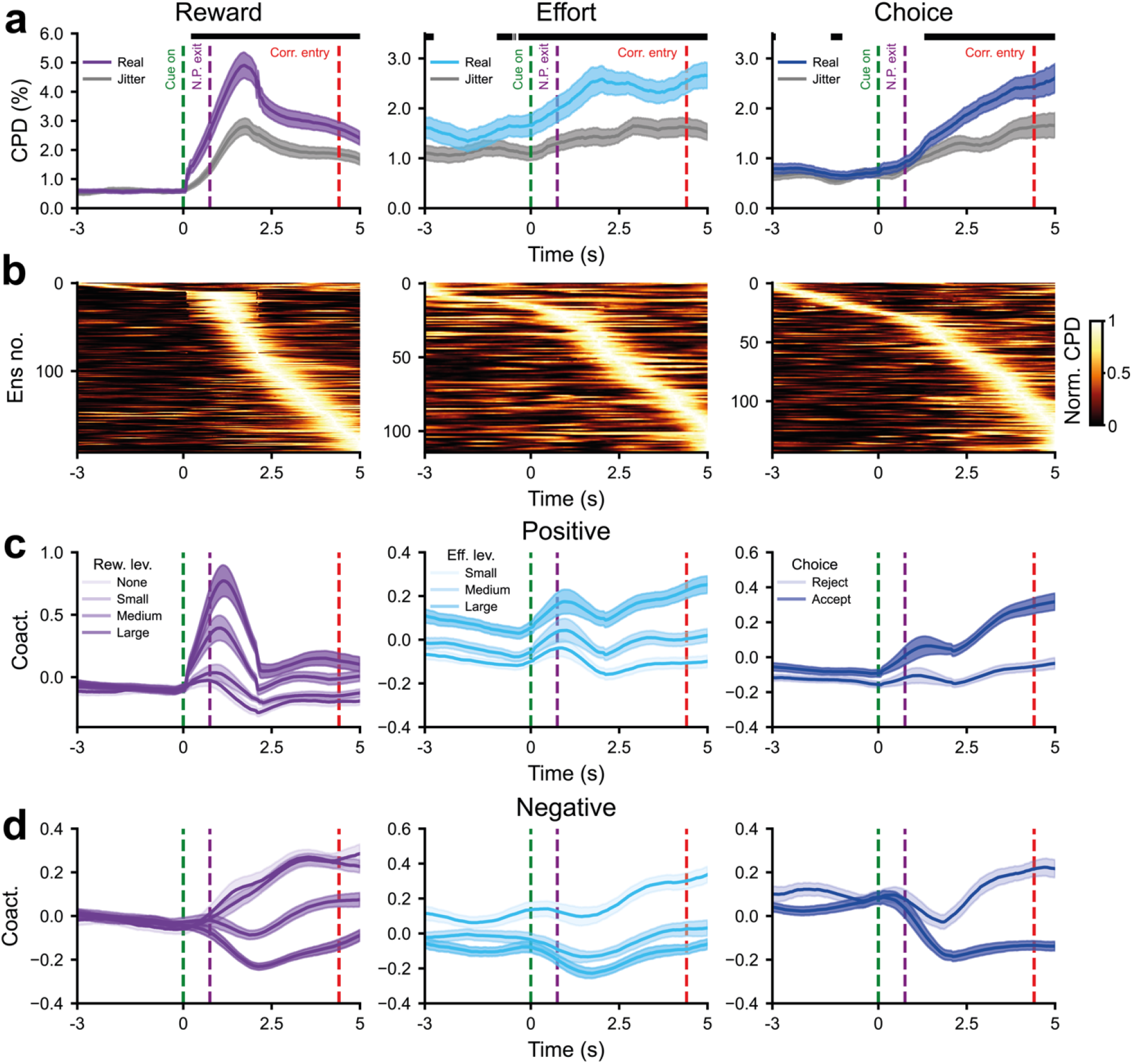
Co-firing assemblies can dynamically encode decision variables on task-relevant timescales. **a**, the CPD for reward, effort and choice regressed onto the time-series of the expression strength of cell assemblies either with (grey) or without (blue) the random jittering of the spike train in the range of -250ms to 250ms for reward, effort and choice coding assemblies (left to right). Time of 0s is the onset of the reward cue and periods of time where the CPD is significantly higher for the unjittered than the jittered assembly pattern expression strength were marked with a blue bar (Wilcoxon signed-rank test, p < .050). The Benjamini-Hochberg procedure was used to control for multiple comparisons. **b**, heat plot where rows represent the normalised coefficient of partial determination (CPD) for reward, effort and choice regressed onto the time-series of the expression strength of cell assemblies for each reward, effort and choice coding assemblies respectively (left to right). CPD was normalised by dividing it by its peak value for each assembly. Time of 0s is the onset of the reward cue. **c** and **d**, the z-scored, smoothed (using a gaussian of std = 2 / √12 𝑠𝑠) coactivity across the different levels of decision variables for positively (**c**) and negatively (**d**) reward, effort and choice (left to right) encoding assemblies. Increasing levels of reward and effort and accept trials were shown in darker colour. Time of 0s is the onset of the reward cue.

Taken together, this shows that assembly co-activity - reflecting temporally co-ordinated activations within and across frontal and basal ganglia structures - dynamically encodes reward and effort as these parameters become known. These are then transformed into an orthogonal choice-related co-activity output reflecting rats’ decisions about whether to pursue or forgo that opportunity.

## Discussion

Cortico-basal ganglia circuits play a key role in choosing actions based on information about prospective costs and benefits, yet the mechanisms behind this have remained elusive. To gain a comprehensive understanding of potential encoding schemes, we developed an “accept-reject” decision paradigm inspired by diet selection foraging ^40^ and previous effort-based decision-making paradigms ^41,42^ and recorded neurons in 5 interconnected regions simultaneously as rats performed this task. Importantly, our task design separated out the initial presentation of the reward offer from subsequent initiation and accept/reject decision, allowing us to dissociate neural activity driven by these factors. ^34,43,44^ Single neurons, most prominently in DMS though also in all recorded regions, signalled information about the reward offer following cue presentation, as well as the current effort block and then the accept/reject decision. Nonetheless, a substantial fraction of all recorded neurons were engaged in a wide range of temporal activity patterns across the task space. Crucially, these same neurons readily formed cell assemblies, dynamically encoding the cost-benefit variables and decisions to act through synchronisation of their spiking activity. Together, these findings suggest a hierarchical organisation of cost-benefit computations, where individual neurons with diverse rate-coding properties are integrated into higher-order, emergent assemblies that can encode decision-related parameters across the circuit at millisecond timescales.

A central question regarding the neural correlates of decision-making is whether decision variables are represented locally or distributed across multiple brain areas ^7,34,44,45^. Lesion studies have shown that ACC and OFC can support dissociable functions during reward-guided choice ^46–48^. If particular computations are supported by particular brain regions, it might be expected that these would be evident in the encoding schemes within those brain regions. The activity of many single neurons in the DMS and STA positively encoded reward in this way, with responses tightly locked to the offer of prospective reward. This finding is in line with the established role of striatal neurons in reward prediction ^49,50^. However, we found that at least 20-30% of neurons in all five structures encoded *each* decision parameter over the entire task window. In fact, even negative reward-encoding neurons in DMS and STA exhibited such temporally distributed activity. Interestingly, while a similar sustained pattern was seen for DMS effort signalling, its encoding scheme also exhibited particular strength around the time the rats initiated a trial. Given that effort levels were uncued and instead fixed over blocks that lasted for minutes, this suggests DMS neurons dynamically recall this information when engaging with the cost-benefit task to guide future behaviour.

Our behavioural results clearly demonstrate that both the current reward value and stored knowledge of the current effort requirement were used to determine the animal’s choice. Neuroimaging studies consistently implicate medial orbitofrontal, dorsal ACC and striatum in signalling the net effort-discounted value of an available reward ^51^ and electrophysiological recordings in primates have suggested that ACC neurons in particular can perform this role ^35,52^. Here, we found that multiplexed combinations of reward and effort were more prevalent in basal ganglia structures, particularly DMS and STA. Notably, this basal ganglia encoding was evident during the cue period prior to any action being initiated. Striatal neurons have previously been shown to integrate reward with other decision variables ^53,54^, but to the best of our knowledge this is the first report of this extending to cost-benefit net value coding.

Having demonstrated that rate-based encoding is generally distributed across the network, we hypothesised that synchronised firing could provide a means through which neurons with diverse tuning properties could be coordinated. We demonstrate that neurons across all the recorded cortico-basal ganglia regions can form cell assemblies, and that these assemblies can encode decision parameters. Cell assemblies have been mostly described in hippocampal neurons during spatial navigation ^25,37^ and during the representation of contextual information ^30^, but to our knowledge, have rarely been described in cortico-basal ganglia networks, and never for key cost-benefit decision variables^27,55^. Here we provide evidence that variability in the strength of cell assembly expression in the cortex, striatum and other basal ganglia structures can individually and conjointly encode multiple levels of prospective reward and effort at timepoints and over timescales relevant to the utilisation of that information to regulate choice behaviour.

Cell assemblies have specific properties that could be as important in value-based learning and decision-making as they are in spatial learning and navigation. Specifically, assemblies promote firing within the membrane time constant (10-30ms) that is optimal to discharge downstream neurons / assemblies and for strengthening synaptic weights between those neurons ^25^. Instead of acting as individual processing units, neurons in cell assemblies, both within and across structures, represent information cooperatively via their synchrony. This synchronous firing can produce a composite downstream affect that cannot be achieved by single neurons alone ^25,26,37^. Therefore, the importance of these findings is that they demonstrate that value-based variables can be represented on a spatiotemporal scale than cannot be achieved by single neurons. These spatiotemporal interactions could promote integration of network-level connectivity akin to that in hippocampal networks, where the key property of the assembly is to simultaneously mobilize enough pyramidal neurons to discharge the target neurons and promote synaptic plasticity between those neurons.

Importantly, our data corroborate a recent demonstration that functional assemblies can span multiple cortico-basal ganglia structures ^27^. While we detected considerably more cross-structural assemblies in absolute numbers than that study, they made up a smaller proportion of assembly patterns overall. The technical challenge of recording large numbers of neurons across structures means that such cross-structural assemblies are probably underrepresented, with actual cell assembly size likely comprised of hundreds of neurons ^56^. Thus, because we sample only a small proportion of assembly members, we likely compute a noisy estimate of the coactivity that arises in the complete cell assembly.

Strikingly, assemblies that encoded a given decision variable were often comprised of single neurons coding for either a different parameter or none of the canonical variables at all (see schematic representation in Fig. 7 for how this can be manifested). This finding supports the idea that such assemblies are *emergent*: the activity of the population is more than a simple sum of the individual members, echoing previous findings in the hippocampus ^30^. This has important implications for future studies of decision-making in these circuits, as it demonstrates that neurons that do not have clear, temporally consistent rate-coding responses (Fig. 2-3) can still play an important role at the population level (Fig. 7). This view is not mutually exclusive to neurons of a specific type having a highly having a specialized role within the network, as striatal neurons appear to do here. Rather, we suggest that higher-level emergent activity constitutes a different level of organisation that facilitates parallel representations and/or computations across neurons and structures in ways that rate-coding alone cannot.

**Figure 7.**
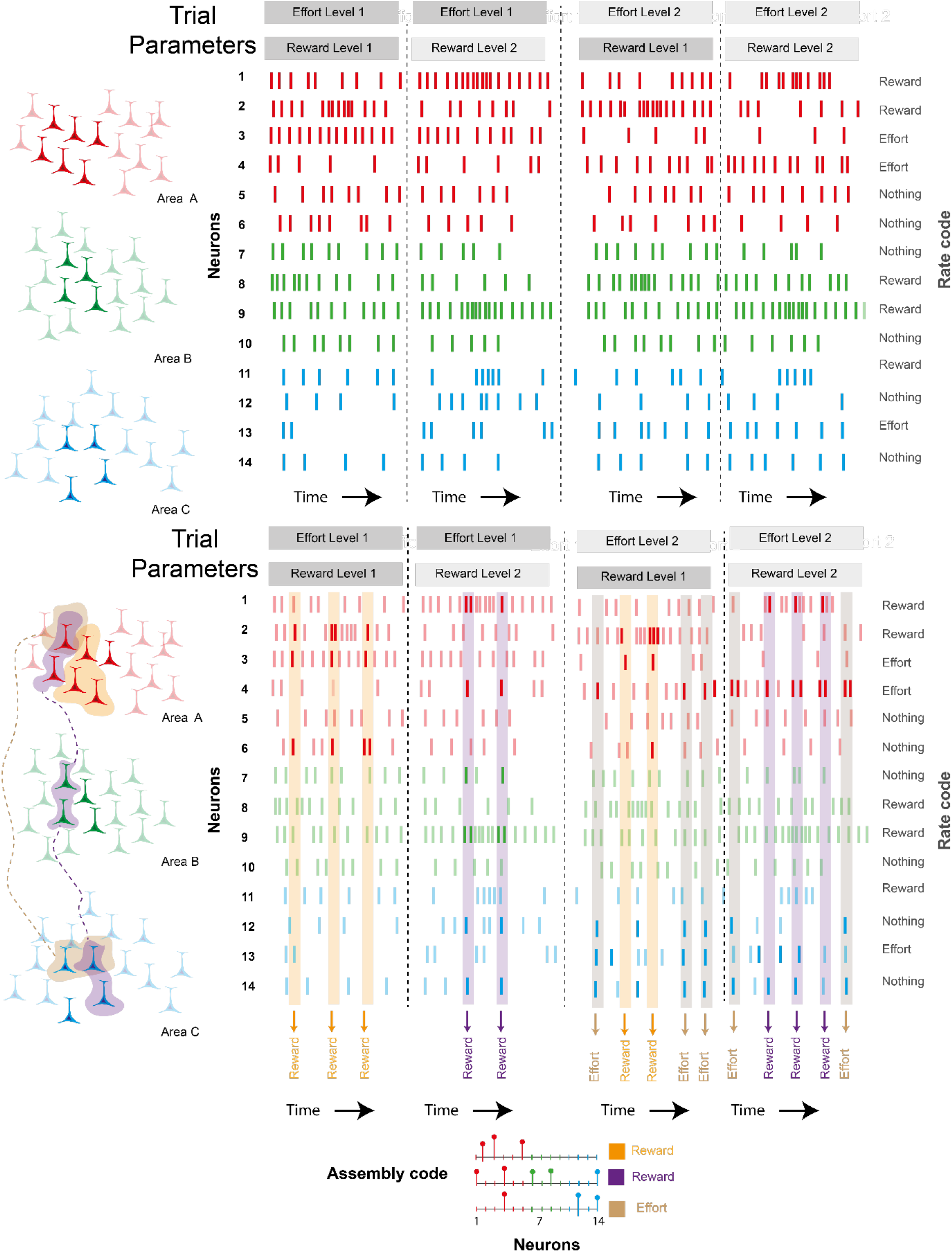
Schematic representation of relationships between detected rate and assembly codes and their possible relationships to the wider network. **a**, Schematic representation of rate responses in 3 populations of neurons (denoted by colour) from different brain areas during trials with different combinations of prospective reward and effort. Neurons can encode the decision parameters and/or transient rate changes that occur consistently in part of the trial (e.g. neurons 1 and 2) and/or global changes in firing rate during specific conditions (e.g. neurons 3 and 4). The recorded neurons (in bold) are assumed to reflect the rate-responses of the wider population (faded) within that structure. **b,** Assembly-coding can occur simultaneously during the rate coding shown in A. Neurons with all types of rate coding (reward, effort, nothing) can contribute to assemblies that encode specific combinations of effort and reward. Spikes that contribute/or a consequence of assemblies can occur inside or outside or rate-coding periods. Assemblies can be formed within and/or across different areas and a given neuron can participate in assemblies encoding different parameters (e.g. neuron 14). The recorded neurons (bold) represent a small part of the “real” assembly which likely comprises many neurons in the wider of population of neurons within and outside the recorded structures.

One key outstanding question is how encoding at the level of single neurons and cell assemblies is related. Few would doubt that the firing rate of a single neuron represents an important level of computation in the brain. Single neuron tuning must ultimately result from integration of synaptic inputs and weights of those inputs to the neuron, which will evolve and consolidate during learning under the influence of neuromodulators such as dopamine. As demonstrated here, such single neuron responses can achieve complex computations, such as the encoding of multiple task variables that occur across different timescales. Nevertheless, while the origin of inputs to a single neuron can arise from disparate and distant brain areas, such computations can only result from the combination of synapses to that neuron.

Cell assemblies, on the other hand, must reflect interactions on a larger spatial scale. Importantly, connections between assembly members need not be direct. DMS neurons had the largest propensity to form assemblies here, but connections between striatal projection neurons are weak ^57^. Striatal assemblies are therefore more likely to be formed from common cortical and thalamic inputs that are the dominant drivers of striatal activity ^24,58^ but require convergence of many neurons to drive reliable output ^59^. Thus, even when an assembly was not defined as cross-structural in our data, it was highly likely to have reflected a much wider pool of connected neurons across structures. This framework shares common ground with the Hopfeldian view that information is represented by coordinated neural activity, conceptualised as an attractor basin in a low dimensional manifold ^60^. Within this approach, the expression strength of each of the identified assemblies can be thought of as dimensions of a latent space describing the neural activity ^61^. This latent space is particularly sensitive to the coactivity of neurons and, importantly, cannot be wholly described by the second-by-second changes in firing rate ^62^. In fact, in the absence of fine temporal coactivity, the position along each dimension of this latent space carries substantially less information about decision relevant variables.

We suggest that the relationship between single neuron rate and assembly codes represent different levels of a hierarchical organisation of information. Assemblies would be higher in the hierarchy in terms of incorporating a larger number of individual neurons and having lower dimensionality. Conversely, assembly formation may be shaped by the tuning curves and afferent/efferent partners of the constituent neurons. Phasically active neurons, such as those in the striatum and subthalamus, may be particularly influential in this regard. Similarly, the expression of assemblies should reinforce the rate-related properties of the individual members by consolidating the synaptic connections. Employing such a nested, hierarchical framework can explain how diverse rate-coding responses can be co-ordinated across brain networks during adaptive behaviour.

## Acknowledgements

We would like to thank T. Akam, M. Rothwell and B. Micklem for methodological and technical assistance and D. Dupret for feedback during the project. This work was supported by The Clarendon Fund and Mary Somerville Clarendon Graduate Scholarship in conjunction with the Department of Experimental Psychology (award SFF1819_CB2_MSD_ 1196514 to OH), the Medical Research Council UK (award MC_UU_00003/6 to AS), the Oxford National Institute for Healthcare Research (NIHR) Biomedical Research Centre to S.G. and the Wellcome Trust (Sir Henry Wellcome Fellowship 209120/Z/17/Z to CGM; 202831/Z/16/Z and 214314/Z/18/Z to MEW).

## Author contributions

Conceptualisation: OH, MEW, AS and SGM; Technical development: OH, RT, TD and CGM; Investigation: OH, BP and IG; Analysis: OH, BP and IG; Resources and funding acquisition: AS and MEW; Writing - original draft: OH, IG, and AS; Writing - Reviewing & Editing: OH, IG, BP, RB, CGM, HC, SGM, MEW, AS; Supervision: AS, MEW, SGM, HC and TD.

## Data Sharing Statement

Upon publication, the data and core analysis code will be made available at https://data.mrc.ox.ac.uk.

## Materials and Methods

### Subjects

All procedures were carried out in accordance with the UK Animals (Scientific Procedures) Act (1986) and its associated guidelines. We used 12 group-housed Lister Hooded naive rats (Charles River, U.K.) weighing 300-390g at the beginning of training. Rats were maintained on a twelve-hour light/dark cycle (lights on at 7 AM). During testing periods, rats were food restricted to a target weight range of 85-90% of their free-feeding weight. Water was available *ad libitum* in their home cages. Following implantation with multielectrode drives, animals were singly housed.

### Behavioural apparatus

The operant box was designed in Fusion360 (Autodesk) and consisted of two 17.6cm diameter custom-designed 3D-printed plastic semi-circular arenas, connected by a 130 cm-long Perspex corridor with walls on either side. The corridor had 6 equidistantly placed infra-red (IR) light-gates to register the rats’ position and 3 equidistant slots for barriers to increase the travel cost between the ends of the corridor. Each arena housed a custom designed 3D-printed plastic nose-poke and reward port, positioned symmetrically on either side. Nose-poke and reward ports contained a infrared light-gate and a white LED house light. Reward ports were connected with 20-mg food pellet dispensers (Med-associates) fitted outside the walls of the arenas using medical-grade silicone tubing. Each reward port housed a moveable floor that was able to obstruct the port. The electronics were controlled by a Teensy 3.6 microcontroller that ran on custom-written firmware.

### Behavioural paradigm

Rats initiated trials by poking and holding in the nose-poke port of the starting arena ("pre-cue"). The starting arena was randomly allocated on the 1st trial. After a pre-cue period (0.3-0.4s), an auditory cue ("offer") was again presented from the distal end of the apparatus to signal the available reward at end of the track from where they initiated. The available reward was pseudo-randomly varied trial-by-trial (4 sizes: nil, low, mid, high, corresponding to zero, one, three or six 20mg sucrose pellets), signaled by a distinct auditory cue. The cue-reward associations were counter-balanced across subjects. Rats were required to maintain nose-poking for another 0.6-0.8s until the onset of a visual Go cue, signaled by a white light LED. Premature exiting of the nose-poke either during the pre-cue or offer period would elicit a 5-s error time-out, indicated by a flashing house-light in the nose-poke, followed by a 5s ITI. If rats exited prematurely during the auditory cue, the same cue would be repeated on the next nose-poke attempt. A grace period of up to 100ms during nose-poking - where nose-poke exits did not count towards an error - allowed for minor adjustments in body position. To accept the offer, rats needed to enter the corridor (cross a light-gate 24cm away from the arena) within a time window; otherwise, that offer was rejected. The cost of pursuing reward was pseudo-randomly varied across blocks of 17-23 trials (3 levels: low, moderate, high), implemented using varying numbers of barriers (0-2 or 1-3 in implanted or non-implanted animals, respectively) in the corridor. The barriers were 0.5 cm thick and were 12 cm and 18 cm tall at their lowest and highest points respectively, with a 45-degree slant mid-way between to generate a more effortful zig-zag of running between the ends of the corridor.

### Behavioural training

Rats were handled daily for a week prior to training. Training consisted of 5 steps in both versions of the task, summarized in table 3.

Step 1 ("magazine training") consisted of rats being introduced into the apparatus with bedding from their home cages and receiving 20 rewards in each food dispenser, which they were free to consume in 20 minutes.

In <u>Step 2</u> ("cue-reward association training") - entry of the starting arena as detected by entry/presence in the arena of latest reward delivery would trigger one of 4 auditory cues, predicting the magnitude of the promised reward in the distal reward port. The offer cue was played until the animal made an entry into the distal reward port. Next, the reward port LED was activated for a 1s pre-reward delay, followed by delivery of the reward corresponding to offer cue. 2.5s after reward delivery, the moveable reward port floor was elevated in that same arena, resulting in the blocking of that port to indicate that reward would not be available there on the next trial. After the rats had reached a criterion of initiating 120 trials in £ 60 minutes, rats were taken to step 3.

<u>In step 3</u> ("nose-poke training"), in v1, rats had to learn to sustain a nose-poke with a jittered 1-1.2s length in order to elicit the offer cue. If rats sustained a 0.3-0.4s-long nose-poke ("pre-cue"), one of 4 auditory cues would be played. Importantly, rats were not free to leave until another 0.6-0.8s later ("offer") once a visual Go cue, signalled by a white light LED in the nose-poke port, had turned on. This was implemented to ensure a movement-free period following offer presentation for electrophysiological analyses. The pre-cue nose-poke requirement was incremented from 0.1s to 0.4s in increments of 25 ms after every successful trial. Once animals had learned to sustain successful pre-cue nose-pokes, the cue nose-poke requirement was incremented from 0.0s to 0.8s in steps of 20 ms after every successful trial. Once animals had learned to complete 120 trials with the full 1.2s nose-poke requirement, jitters of 0.1s ("pre-cue") and 0.2s ("offer") were introduced. To allow for small changes in body posture that may have caused the nose-poke sensor to stop registering an entry, a 0.1s grace period was introduced which did not result in a nose-poke error. Importantly, if a nose-poke was interrupted for more than 0.1s (the grace period) during the "offer" window, the same offer cue would be repeated during the next trial. This was not an intentional feature, but rather the side-effect of using a pre-determined sequence of offers. Once rats had reached a criterion of initiating 120 trials within 60 minutes, they were taken on to step 4.

<u>In step 4</u> ("barrier training") rats needed to nose-poke to elicit offer cues as outlined previously, but now needed to additionally learn to scale barriers when traversing the corridor between the arenas. Barriers were added in one by one in blocks of 40 trials, starting from 1 barrier and going up to a maximum of 3 barriers. Once rats had successfully managed to complete 120 trials within 60 mins, they were taken on to step 5.

<u>In step 5</u> ("final training for non-implanted animals") *r*ats were required to either accept or reject a distal reward offer available at the other side of the corridor (fig. 2). After the onset of the Go cue, to accept the offer, rats needed to enter the corridor within a 3s time window as determined by a light-gate placed near the border of the arena; otherwise, that offer is rejected. The effort cost which was fixed across blocks of ∼20 trials (3 levels: low – 1 barrier; moderate – 2 barriers; high – 3 barriers; table 3). Upon corridor entry, they were allowed a further 5s to traverse the corridor and reach the distal reward port. To ensure average reward rate was minimally affected by rats’ accept/reject choices, ITIs were adjusted daily on ‘accept’ trials by subtracting the previous day’s cohort median run time.

<u>In step</u> 5 ("post-surgery re-training"), the corridor entry and traverse times were increased to 15s and 10s respectively post-surgery to account for the additional constraints on implanted rats with a recording tether. Additionally, to avoid collisions between the implant and the barriers, the number of barriers per each level of effort was reduced by 1. Rats were considered ready for the recording experiment once they managed to complete 120 trials within 60 minutes.

### Outcome devaluation experiment

Following standard testing, outcome devaluation was performed on a subset of rats (n=6, first cohort) in a within-subjects regular Latin square design. In devaluation sessions, subjects were placed in a solitary "feeding" cage with *ad libitum* lab chow for 1h prior to the start of the testing session. Rats were placed in a solitary "control" cage for 1h on control sessions. The outcome devaluation lasted 3 days altogether, out of which days 1 and 3 were testing days and day 2 was a baseline training day.

### Multielectrode drive

For single unit recordings we used a bespoke driveable multielectrode device weighing ∼20g. The implant was comprised of a lid which housed a 128-pin connector designed for Intan 64-channel head-stages (Intan Technology), a plastic corpus that housed the electrode movement mechanisms, glass tubing and a base. All components were assembled in house. For assembly, the elements of the corpus were first 3D printed and post-processed with UV radiation and propanol. Second, holes were drilled in the corpus to allow threading of electrode wires and the driving mechanisms. Third, driving support mechanisms were created using metal rods, plastic shuttles and screws. Fourth, glass tubing was introduced into the driving mechanisms as electrode supports. Fifth, strands of 50-micron diameter tungsten wire (California Fine Wire) were twisted and heated to create 4- and 6-strand electrodes (tetrodes and hextrodes, respectively). Sixth, these were threaded through the glass tubing and the rest of the implant corpus. Seventh, ground wire stubs (connected to ground wires on the animal during surgery, see below) were introduced into the implant using a dedicated set of stationary holes. Finally, the implant was sealed and the electrodes cut to length by hand.

Three of the subjects (gh001, gh009, gh010) reported here received an implant that targeted all 5 regions, whereas one subject (oh011) received a modified drive without the VP electrodes (table 1). Electrodes in the MO/VO, ACC, and VP were arranged as a row along one sagittal plane. DMS electrodes were as rows along two sagittal planes and STA electrodes were arranged in triangular fashion around a centre point. The ML and AP positions of each electrode in the drive were planned around a centre-point co-ordinate obtained as an average between readings in two different versions of a commonly used rat brain atlas ^1,2^ that visually offered the most exposure in each structure, while also avoiding the ventricles (Fig. S2a). Relative electrode positions in the AP and ML dimensions were planned around centroid co-ordinates.

**Table 1.**
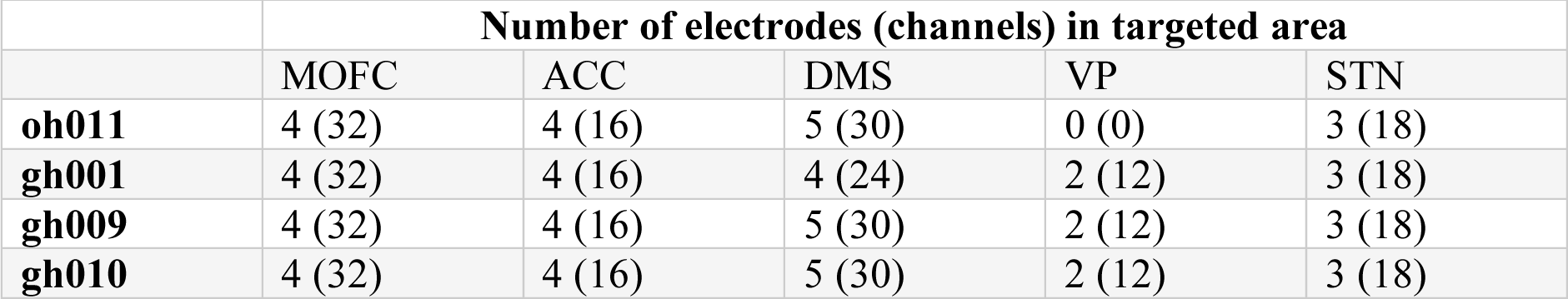
Overview of the number of electrodes and channels in each subject and targeted recording area.

### Implantation of multielectrode drives for recordings in behaving rats

Electrode implantations were performed in a subset of expert animals (n=4; table 1). Surgeries were performed under isoflurane anaesthesia (5% for induction, 1.5-2.5% maintenance) and oxygen (2L/min). Subcutaneous local anaesthetic (Marcaine, 2mg/kg, 2.5 mg/mL) and a non-steroidal anti-inflammatory drug (Metacam, 1mg/kg, 5mg/mL) were administered at the beginning of each surgery. Post-operative opioid analgesia (Vetergersic, 0.3 mg/mL, 0.03 mg/kg) was administered subcutaneously for three consecutive post-operative days.

Craniotomies were created above each of the targeted areas and *dura mater* thoroughly removed from the visible brain surface. The drive was aligned on the animals’ heads by using the centre-point of the STA electrode bundle as a reference. Next, the drive was lowered to target depth on a stereotaxic frame (David Kopf Instruments) based on the DV reading of the STA electrode bundle from brain surface. The implant was then secured to skull-attached screws using adhesive dental cement and dental acrylic. Recordings were referenced to two supra-cerebellar screws which were connected to the implant during surgery. Neuronal recordings began at least 2 weeks after surgery. Electrode placement was confirmed using histology.

### Histological data analysis

To verify electrode placements, after the experiments were completed, animals were deeply anaesthetised with isoflurane (5%) and intraperitoneal pentobarbital (3 mL, Pentoject, 200 mg/mL and perfused trans-cardially with phosphate buffered saline (PBS) followed by fixative solution (paraformaldehyde dissolved in PBS, 4%, wt/vol). For post-mortem fixation, the sectioned heads were further submerged in fixative for another 24h, following which brains were extracted and washed 3 times in PBS and stored in a PBS solution containing 0.02% sodium azide until sectioning. Brains were sectioned at 0.1mm thickness using a vibratome (Leica), immunohistologically stained and mounted in Vectashield. Unfiltered "bright-field" epifluorescence microscope images were using 5x magnification and Zen Blue Pro (Zeiss).

### Trode localisation and histological verification

To estimate all the possible recorded areas and exclude cells yields from neighbouring regions, we (1) estimated each identifiable electrode tract’s DV end-depth from our epifluorescence imaging data, (2) averaged this across all identifiable electrode tracts to get an average end-depth, (3) estimated each day’s targeted structure based on a log of putative lowering distances averaged across electrodes within a region. We did not restrict the cell yield for the STA based on histological data. For other structures we restricted our cell yields to only the MO or VO for MO/VO, Cg1 and Cg2 for the ACC, CPu for the DMS; and VP only for the VP.

### Electrophysiology data acquisition and processing

Wideband and accelerometer data were recorded, amplified and digitised on 2 tethered Intan 64-channel headstages at 20,000 Hz using an Intan acquisition board (Intan Technologies). No hardware filtering was applied during recording. Files were saved in binary format. Each electrode was lowered by at least 125 mm up to 2h ahead of each recording session to ensure identification of unique units every day.

### Behavioural data analysis

All data were analysed in Python. A binomial mixed effects model using the pymer4 library ^3^ was used to regress the following fixed effects onto trial-by-trial choice:

- *reward*, coded as 0, 1, 3 or 6
- *effort*, coded as 1, 2, or 3 (0, 1, or 2 in implanted animals)
- *reward x effort interaction*, coded as the multiplication of those vectors
- *reward count*, coded as the number of pellets earned since the start of the session.

To safeguard against type I errors, we aimed to fit a maximal random effects structure whenever possible by modelling each predictor as a random, in addition to a fixed, slope ^4^. To account for day-by-day and subject-by-subject covariation in the data, random slopes were paired with a random intercept for either the testing day or the subject ID. Whenever model fitting failed to converge, random effects that fit close to zero variance were removed to simplify the random effects’ structure. The resulting decision model for the non-implanted cohort had the following formula:

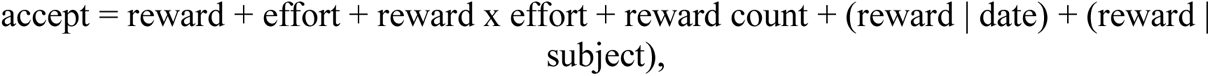

while the implanted rat decision model was formulated as follows:

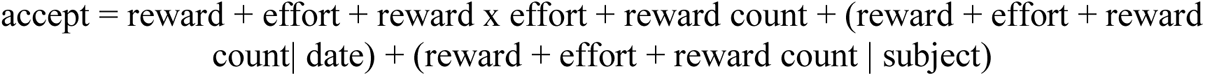

For the outcome devaluation experiment, we added pre-feeding and its interaction with reward and effort as an additional predictor, resulting in the following formula:

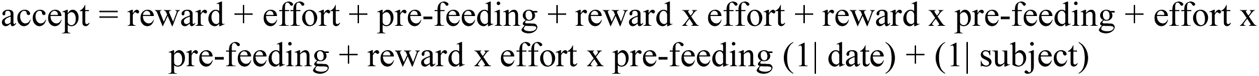

All predictors were standardised to facilitate side-by-side comparison. P-values were obtained using the likelihood ratio test.

### Firing rate modulation during task execution

For visualising task event-locked firing rates (Fig. 2b), spiking counts were quantified in 10ms non-overlapping bins sampled in 10 ms increments -400ms to +400ms around the initiation of the nose-poke (termed "Pre-Cue"), -400ms to +400ms around the onset of the offer cue (termed "Offer"), -500ms to +500ms around the onset of the Go cue (termed "Go Cue") and -500ms pre-to +1000ms following nose-poke exit (termed "Action"). Data were subsequently averaged across trials of the same magnitude of offered reward, the required effort and prospective decision, each time averaging across the other two variables. Peri-stimulus time histogram (PSTH) data were smoothed with a gaussian filter of standard deviation 5.0 for display purposes.

To facilitate the alignment of data on the task (Fig. 3; Fig S3-4), time on trial was transformed into a uniform bin space by dividing the Pre-Cue epoch (-400 ms before Pre-Cue up to Offer) into 16 bins, the Offer epoch (from Offer up to Go Cue) into 16 bins, the Go Cue epoch (from Go Cue up to Action) into 13 bins and the Action epoch (from Action to corridor entry on accepted trials or each animal’s daily median accepted trial corridor entry latency on rejected trials) into 18 bins. Firing rates were quantified in 50 ms bins centered around each such bin and z-scored relative to a concatenated series of ITI data from the ongoing effort block.

### Individual neuron regression analyses

General linear models (GLMs) were employed to quantify the influence of reward, effort and prospective decision on z-scored individual unit firing rates. For this, the loading (b) of each variable on z-scored firing rates were obtained using the closed-form least-squares solution:

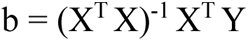

where X is a design matrix of shape [trials x variables] and Y is a vector of z-scored firing rates of shape [trials x 1] from one recording day. The 4 regressors used in the design matrix X were constructed as follows:

- Intercept: a vector of ones
- Reward: a vector containing the values 1, 3 or 6 corresponding to small, medium or large reward trials.
- Effort: a vector containing the values 1, 2 or 3 corresponding to low, moderate or high effort trials.
- Decision: a vector comprised of the values -0.5 and +0.5 corresponding to rejected or accepted trials.

All regressors except the intercept were z-scored to themselves before fitting the models and obtaining coefficient loadings. From these GLMs, coefficients of partial determination (CPD), reflecting the unique variance explained by variable, were obtained as follows:

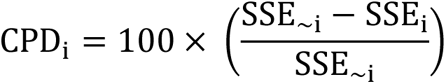

where SSE_∼i_ and SSE_i_ represent the sum of squared errors of models that, respectively, do not or do include the regressor of interest. Using CPD ensured that the encoding strength for a given variable was not secondary to another (e.g. higher firing rate/encoding on accept trials due to more of these trials having higher prospective reward). To identify significant coding, permutation testing was used. For each neuron and decision variable, 1,000 shuffled datasets were generated by randomly shuffling the trial indices within each recording day. Such shuffling preserves potential autocorrelations between task variables and firing rates and is thus an appropriate estimate of the chance-level outcome. Individual neuron regression coefficient p-values were calculated as the fraction of (1,000) permuted CPD values which exceeded a neuron’s non-permuted CPD.

Significant neurons (Fig. 2b; Fig 3c: Fig. S3-4) were identified on the basis of CPD p-values less than 0.05 across in one single time bin abutting the onset of the offer cue and entry into the corridor (on accepted trials) or the session median corridor entry latency (on rejected trials). Individual neuron examples in Fig. 2b were selected if they met this criterion and displayed some consistency in the discrimination response over time based on visual assessment. Instantaneous population CPD p-values (Fig. 3a) were calculated as the fraction of permutations for which the (1,000) shuffled population average CPDs were larger than the non-permuted population mean CPD. The Benjamini-Hochberg (BH) procedure was applied to correct for false discovery rate across tested time points.

The proportions of significantly coding neurons between regions were compared pair-wise between areas using the z-test of proportions with BH correction for the number of unique comparisons.

### Quantification of CPD phasicness (fig. 3a insets)

To characterise the degree to which individual unit CPDs magnitudes were temporally localised vs distributed across the trial timeline, we calculated *phasicness.* This is related to *sparsity* as defined by previous studies ^5,6^ and indexed the extent to which the CPD time series for a population of neurons was dominated by a small number of time points. For each neuron in each area, it was defined as follows:

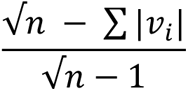

where n is the length of the CPD vector (i.e., the number of time points) and v_i_ is the CPD vector’s i^th^ element. For this analysis, the input CPD time series was a unit vector where the sum of the squares of all elements added up to one. Phasicness distributions were compared between brain areas using the two-sided Mann-Whitney U test with BH correction for the number of pairwise comparisons. All Mann-Whitney U test results reported in this work as expressed as a Z-score obtained as:

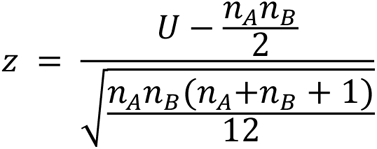

where U is the Mann-Whitney U test statistic and n_A_ and n_B_ represent the sample sizes in the two comparison groups.

### Quantification of latency to peak coding response and phasic responding properties (fig. 3a, lower panel)

To quantify the latency to peak coding response, we calculated the time bin of maximal CPD magnitude for each neuron and plotted the distribution across all neurons as violin plots using the python Seaborn library. To evaluate whether these peak CPD latencies tended to co-occur close-by in time (phasic responding) more than would expected by chance, we compared these values with a null-distribution formed by bootstrapped sampling of all observed unique latency values 1,000 times with replacement. The null and real distributions were compared using the Mann-Whitney U test and the false-discovery rate corrected for by using the BH correction.

### Co-activity analysis

Assembly patterns were determined using the combination of principal component analysis and independent component analysis (PCA-ICA) as detailed earlier ^7,8^. This analysis was conducted using neurons from all structures simultaneously. This allowed for the unsupervised extraction of assembly patterns which could be contained entirely within an anatomical structure or could span multiple brain regions. The spike trains for all units during time periods where the animal was actively engaged with the task were binned at 25ms, z-scored and concatenated into a *n* x *B* matrix, where *n* was the number of neurons and *B* was the number of 25ms bins. The element of this matrix at position (i, b) was the z-scored spike count of neuron i in bin b.

PCA was then applied to the z-scored spike count matrix. The number of significant patterns embedded in the z-scored spike counts was determined using the Marcenko-Pastur law. This is an analytical result from random matrix theory, which bounds the magnitude of the eigenvalues from PCA that can occur due to chance. Principal components with eigenvalues above the upper limit of the bound set by the Marcenko-Pastur law were classified as significant. Principal components with an eigenvalue greater than this threshold captured more of the correlation which existed between the binned firing of neurons than should occur if all neurons spiked independently of one and other ^5^.

The z-scored spike count matrix was then projected onto the significant principal components and ICA was applied to the resulting matrix. The output of ICA was an unmixing matrix. This unmixing matrix was then projected back onto the original basis, where each dimension represents the binned z-scored spike count of an individual neuron. The columns of the resulting matrix represent assembly patterns. These assembly patterns contain a weight for each neuron. The greater this weight is, the greater the contribution the neuron makes to the identified coactivity pattern ^5^.

There are several advantages to using the combination of PCA and ICA as opposed to using either technique alone ^5,7,8^. For instance, PCA has been found to group several assembly patterns into the first principal component. PCA also artificially constrains identified assembly patterns to be orthogonal to each other, meaning that a high weighting of a neuron in one assembly pattern limits the weighting of that neuron is subsequently identified patterns. Finally, PCA is only based on pairwise correlations and cannot identify higher order correlations. However, if ICA was applied directly to the z-scored binned spike count matrix, one could identify the same number of assembly patterns as there are neurons. This could result in the extraction of spurious patterns. To address these caveats, ICA is applied to the z-scored binned spike count matrix following dimensionality reduction using PCA.

In order to determine the expression of assembly patterns across time, the outer product of each assembly pattern was determined ^5^.The expression strength (R) of assembly pattern k can be determined as follows:

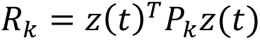

where P_k_ is the outer product of assembly pattern k with elements in the main diagonal set to 0, and z(t) is a column vector containing the smoothed z-scored firing rate of neurons at time t. The main diagonal of 𝑃𝑃_𝑘𝑘_ must be set to zero to ensure that assembly pattern expression strength cannot be driven solely by isolated increases in the firing rate of individual units. The instantaneous firing rate of neurons in 𝑧𝑧(𝑡𝑡) were computed by convolving the spike train of neuron with a gaussian kernel and then z-scoring the resulting timeseries for each neuron. The standard deviation of the gaussian kernel was set to 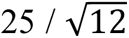 ms to match the 25ms bin used for PCA-ICA ^9^. Convolution with a gaussian kernel allows for greater temporal resolution when compared to the 25ms bins which were used to define the assembly patterns via PCA-ICA.

Linear and logistic regression was conducted using either the python module statsmodel or in custom written pytorch functions. The general strategy utilized in the linear regressions was to take an epoch in each trial, determine the value of variables such as the spike rate or the mean coactivity in this epoch and use this to predict reward, effort or choice outcome across trials, or vice versa.

For identifying coding ensembles, the mean expression strength of an ensemble was calculated from an interval starting with offer cue onset and ending with either corridor entry or, in instances where the animal did not enter the corridor, the median duration taken for corridor entry on that recording session. The mean expression strength was then fitted as the dependent variable in a linear regression with reward, effort and choice outcome as independent regressors. Significance was determined by comparing the CPD of reward, effort and choice to that of the 95𝑡ℎ percentile of the null distribution where reward and effort levels or choice outcomes were shuffled across trials. After identifying ensembles with significant modulation by the level of reward, effort or choice, we probed the time course of this modulation. This was achieved by fitting regression models with reward, effort and choice outcome as independent variables to the smoothed assembly pattern expression strength at different time points. Smoothing was achieved by convolution with a Gaussian with a standard deviation of 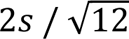. The CPD for reward, effort and choice was compared to that of linear models fitted to the assembly pattern expression strength in jittered spike trains.

Data Accessibility Statement: Data is not available at this Preprint stage due to ongoing work.

## References

1 Adam, R. et al. Dopamine reverses reward insensitivity in apathy following globus pallidus lesions. Cortex 49, 1292–1303 (2013). 10.1016/j.cortex.2012.04.013

2 Hauber, W. & Sommer, S. Prefrontostriatal circuitry regulates effort-related decision making. Cereb Cortex 19, 2240–2247 (2009). 10.1093/cercor/bhn241

3 Levy, R. & Dubois, B. Apathy and the functional anatomy of the prefrontal cortex-basal ganglia circuits. Cereb Cortex 16, 916–928 (2006). 10.1093/cercor/bhj043

4 Manohar, S. G. & Husain, M. Human ventromedial prefrontal lesions alter incentivisation by reward. Cortex 76, 104–120 (2016). 10.1016/j.cortex.2016.01.005

5 Munster, A. & Hauber, W. Medial Orbitofrontal Cortex Mediates Effort-related Responding in Rats. Cereb Cortex 28, 4379–4389 (2018). 10.1093/cercor/bhx293

6 Pennartz, C. M. et al. Corticostriatal Interactions during Learning, Memory Processing, and Decision Making. J Neurosci 29, 12831–12838 (2009). 10.1523/JNEUROSCI.3177-09.2009

7 Rushworth, M. F., Noonan, M. P., Boorman, E. D., Walton, M. E. & Behrens, T. E. Frontal cortex and reward-guided learning and decision-making. Neuron 70, 1054–1069 (2011). 10.1016/j.neuron.2011.05.014

8 Uslaner, J. M., Yang, P. & Robinson, T. E. Subthalamic nucleus lesions enhance the psychomotor-activating, incentive motivational, and neurobiological effects of cocaine. J Neurosci 25, 8407–8415 (2005). 10.1523/JNEUROSCI.1910-05.2005

9 Walton, M. E., Bannerman, D. M., Alterescu, K. & Rushworth, M. F. Functional specialization within medial frontal cortex of the anterior cingulate for evaluating effort-related decisions. J Neurosci 23, 6475–6479 (2003). 10.1523/JNEUROSCI.23-16-06475.2003

10 Abitbol, R. et al. Neural mechanisms underlying contextual dependency of subjective values: converging evidence from monkeys and humans. J Neurosci 35, 2308–2320 (2015). 10.1523/JNEUROSCI.1878-14.2015

11 Balewski, Z. Z., Elston, T. W., Knudsen, E. B. & Wallis, J. D. Value dynamics affect choice preparation during decision-making. Nat Neurosci 26, 1575–1583 (2023). 10.1038/s41593-023-01407-3

12 Cowen, S. L., Davis, G. A. & Nitz, D. A. Anterior cingulate neurons in the rat map anticipated effort and reward to their associated action sequences. J Neurophysiol 107, 2393–2407 (2012). 10.1152/jn.01012.2011

13 Diehl, G. W. & Redish, A. D. Differential processing of decision information in subregions of rodent medial prefrontal cortex. Elife 12 (2023). 10.7554/eLife.82833

14 Hillman, K. L. & Bilkey, D. K. Neurons in the rat anterior cingulate cortex dynamically encode cost-benefit in a spatial decision-making task. J Neurosci 30, 7705–7713 (2010). 10.1523/JNEUROSCI.1273-10.2010

15 Lopatina, N. et al. Medial Orbitofrontal Neurons Preferentially Signal Cues Predicting Changes in Reward during Unblocking. J Neurosci 36, 8416–8424 (2016). 10.1523/JNEUROSCI.1101-16.2016

16 Nakamura, K., Santos, G. S., Matsuzaki, R. & Nakahara, H. Differential reward coding in the subdivisions of the primate caudate during an oculomotor task. J Neurosci 32, 15963–15982 (2012). 10.1523/JNEUROSCI.1518-12.2012

17 Nougaret, S., Baunez, C. & Ravel, S. Neurons in the Monkey’s Subthalamic Nucleus Differentially Encode Motivation and Effort. J Neurosci 42, 2539–2551 (2022). 10.1523/JNEUROSCI.0281-21.2021

18 Porter, B. S., Hillman, K. L. & Bilkey, D. K. Anterior cingulate cortex encoding of effortful behavior. J Neurophysiol 121, 701–714 (2019). 10.1152/jn.00654.2018

19 San-Galli, A., Varazzani, C., Abitbol, R., Pessiglione, M. & Bouret, S. Primate Ventromedial Prefrontal Cortex Neurons Continuously Encode the Willingness to Engage in Reward-Directed Behavior. Cereb Cortex 28, 73–89 (2018). 10.1093/cercor/bhw351

20 Tachibana, Y. & Hikosaka, O. The primate ventral pallidum encodes expected reward value and regulates motor action. Neuron 76, 826–837 (2012). 10.1016/j.neuron.2012.09.030

21 Kable, J. W. & Glimcher, P. W. The neurobiology of decision: consensus and controversy. Neuron 63, 733–745 (2009). 10.1016/j.neuron.2009.09.003

22 Padoa-Schioppa, C. Neurobiology of economic choice: a good-based model. Annu Rev Neurosci 34, 333–359 (2011). 10.1146/annurev-neuro-061010-113648

23 Rangel, A. & Hare, T. Neural computations associated with goal-directed choice. Curr Opin Neurobiol 20, 262–270 (2010). 10.1016/j.conb.2010.03.001

24 Peters, A. J., Fabre, J. M. J., Steinmetz, N. A., Harris, K. D. & Carandini, M. Striatal activity topographically reflects cortical activity. Nature 591, 420–425 (2021). 10.1038/s41586-020-03166-8

25 Buzsaki, G. Neural syntax: cell assemblies, synapsembles, and readers. Neuron 68, 362–385 (2010). 10.1016/j.neuron.2010.09.023

26 Hebb, D. O. The organization of behavior; a neuropsychological theory. (Wiley, 1949).

27 Oberto, V. J. et al. Distributed cell assemblies spanning prefrontal cortex and striatum. Curr Biol 32, 1–13 e16 (2022). 10.1016/j.cub.2021.10.007

28 van de Ven, G. M., Trouche, S., McNamara, C. G., Allen, K. & Dupret, D. Hippocampal Offline Reactivation Consolidates Recently Formed Cell Assembly Patterns during Sharp Wave-Ripples. Neuron 92, 968–974 (2016). 10.1016/j.neuron.2016.10.020

29 Wilson, M. A. & McNaughton, B. L. Reactivation of hippocampal ensemble memories during sleep. Science 265, 676–679 (1994). 10.1126/science.8036517

30 El-Gaby, M. et al. An emergent neural coactivity code for dynamic memory. Nat Neurosci 24, 694–704 (2021). 10.1038/s41593-021-00820-w

31 Pessiglione, M., Vinckier, F., Bouret, S., Daunizeau, J. & Le Bouc, R. Why not try harder? Computational approach to motivation deficits in neuro-psychiatric diseases. Brain 141, 629–650 (2018). 10.1093/brain/awx278

32 Paxinos, G. & Watson, C. The rat brain in stereotaxic coordinates. (Elsevier, 2006).

33 Shamash, P., Carandini, M., Harris, K. D. & Steinmetz, N. A. (2018). 10.1101/447995

34 Cisek, P. Making decisions through a distributed consensus. Curr Opin Neurobiol 22, 927–936 (2012). 10.1016/j.conb.2012.05.007

35 Kennerley, S. W. & Wallis, J. D. Evaluating choices by single neurons in the frontal lobe: outcome value encoded across multiple decision variables. Eur J Neurosci 29, 2061–2073 (2009). 10.1111/j.1460-9568.2009.06743.x

36 Roesch, M. R. & Olson, C. R. Neuronal activity related to reward value and motivation in primate frontal cortex. Science 304, 307–310 (2004). 10.1126/science.1093223

37 Harris, K. D., Csicsvari, J., Hirase, H., Dragoi, G. & Buzsaki, G. Organization of cell assemblies in the hippocampus. Nature 424, 552–556 (2003). 10.1038/nature01834

38 Lopes-dos-Santos, V., Conde-Ocazionez, S., Nicolelis, M. A., Ribeiro, S. T. & Tort, A. B. Neuronal assembly detection and cell membership specification by principal component analysis. PLoS One 6, e20996 (2011). 10.1371/journal.pone.0020996

39 Lopes-dos-Santos, V., Ribeiro, S. & Tort, A. B. Detecting cell assemblies in large neuronal populations. J Neurosci Methods 220, 149–166 (2013). 10.1016/j.jneumeth.2013.04.010

40 Stephens, D. W. a. K., J.R. . Foraging Theory., (Princeton University Press, 1986).

41 Le Heron, C., Apps, M. A. J. & Husain, M. The anatomy of apathy: A neurocognitive framework for amotivated behaviour. Neuropsychologia 118, 54–67 (2018). 10.1016/j.neuropsychologia.2017.07.003

42 Muller, T., Klein-Flugge, M. C., Manohar, S. G., Husain, M. & Apps, M. A. J. Neural and computational mechanisms of momentary fatigue and persistence in effort-based choice. Nat Commun 12, 4593 (2021). 10.1038/s41467-021-24927-7

43 Christensen, A. J., Ott, T. & Kepecs, A. Cognition and the single neuron: How cell types construct the dynamic computations of frontal cortex. Curr Opin Neurobiol 77, 102630 (2022). 10.1016/j.conb.2022.102630

44 Hunt, L. T. & Hayden, B. Y. A distributed, hierarchical and recurrent framework for reward-based choice. Nat Rev Neurosci 18, 172–182 (2017). 10.1038/nrn.2017.7

45 Yoo, S. B. M. & Hayden, B. Y. Economic Choice as an Untangling of Options into Actions. Neuron 99, 434–447 (2018). 10.1016/j.neuron.2018.06.038

46 Camille, N., Tsuchida, A. & Fellows, L. K. Double dissociation of stimulus-value and action-value learning in humans with orbitofrontal or anterior cingulate cortex damage. J Neurosci 31, 15048–15052 (2011). 10.1523/JNEUROSCI.3164-11.2011

47 Rudebeck, P. H. et al. Frontal cortex subregions play distinct roles in choices between actions and stimuli. J Neurosci 28, 13775–13785 (2008). 10.1523/JNEUROSCI.3541-08.2008

48 Rudebeck, P. H., Walton, M. E., Smyth, A. N., Bannerman, D. M. & Rushworth, M. F. Separate neural pathways process different decision costs. Nat Neurosci 9, 1161–1168 (2006). 10.1038/nn1756

49 Oyama, K., Hernadi, I., Iijima, T. & Tsutsui, K. Reward prediction error coding in dorsal striatal neurons. J Neurosci 30, 11447–11457 (2010). 10.1523/JNEUROSCI.1719-10.2010

50 Shin, E. J. et al. Robust and distributed neural representation of action values. Elife 10 (2021). 10.7554/eLife.53045

51 Lopez-Gamundi, P. et al. The neural basis of effort valuation: A meta-analysis of functional magnetic resonance imaging studies. Neurosci Biobehav Rev 131, 1275–1287 (2021). 10.1016/j.neubiorev.2021.10.024

52 Kennerley, S. W., Dahmubed, A. F., Lara, A. H. & Wallis, J. D. Neurons in the frontal lobe encode the value of multiple decision variables. J Cogn Neurosci 21, 1162–1178 (2009). 10.1162/jocn.2009.21100

53 Bloem, B. et al. Multiplexed action-outcome representation by striatal striosome-matrix compartments detected with a mouse cost-benefit foraging task. Nat Commun 13, 1541 (2022). 10.1038/s41467-022-28983-5

54 Rueda-Orozco, P. E. & Robbe, D. The striatum multiplexes contextual and kinematic information to constrain motor habits execution. Nat Neurosci 18, 453–460 (2015). 10.1038/nn.3924

55 Trouche, S. et al. A Hippocampus-Accumbens Tripartite Neuronal Motif Guides Appetitive Memory in Space. Cell 176, 1393–1406 e1316 (2019). 10.1016/j.cell.2018.12.037

56 Ainsworth, M. et al. Rates and rhythms: a synergistic view of frequency and temporal coding in neuronal networks. Neuron 75, 572–583 (2012). 10.1016/j.neuron.2012.08.004

57 Koos, T., Tepper, J. M. & Wilson, C. J. Comparison of IPSCs evoked by spiny and fast-spiking neurons in the neostriatum. J Neurosci 24, 7916–7922 (2004). 10.1523/JNEUROSCI.2163-04.2004

58 Hunnicutt, B. J. et al. A comprehensive excitatory input map of the striatum reveals novel functional organization. Elife 5 (2016). 10.7554/eLife.19103

59 Ramanathan, S., Hanley, J. J., Deniau, J. M. & Bolam, J. P. Synaptic convergence of motor and somatosensory cortical afferents onto GABAergic interneurons in the rat striatum. J Neurosci 22, 8158–8169 (2002). 10.1523/JNEUROSCI.22-18-08158.2002

60 Barack, D. L. & Krakauer, J. W. Two views on the cognitive brain. Nat Rev Neurosci 22, 359–371 (2021). 10.1038/s41583-021-00448-6

61 Langdon, C., Genkin, M. & Engel, T. A. A unifying perspective on neural manifolds and circuits for cognition. Nat Rev Neurosci 24, 363–377 (2023). 10.1038/s41583-023-00693-x

62 Stringer, C. et al. Spontaneous behaviors drive multidimensional, brainwide activity. Science 364, 255 (2019). 10.1126/science.aav7893

## References

1 Paxinos, G. & Watson, C. The Rat Brain Atlas in Stereotaxic Coordinates. 4th edn, (Elsevier, 1998).

2 Paxinos, G. & Watson, C. The Rat Brain Atlas. 6th edn, (Elsevier, 2006).

3 Jolly, E. Pymer4: Connecting R and Python for Linear Mixed Modeling. Journal of Open Source Software 3 (2018). 10.21105/joss.00862

4 Barr, D. J., Levy, R., Scheepers, C. & Tily, H. J. Random effects structure for confirmatory hypothesis testing: Keep it maximal. J Mem Lang 68 (2013). 10.1016/j.jml.2012.11.001

5 van de Ven, G. M., Trouche, S., McNamara, C. G., Allen, K. & Dupret, D. Hippocampal Offline Reactivation Consolidates Recently Formed Cell Assembly Patterns during Sharp Wave-Ripples. Neuron 92, 968–974 (2016). 10.1016/j.neuron.2016.10.020

6 McHugh, S. B. et al. Adult-born dentate granule cells promote hippocampal population sparsity. Nat Neurosci 25, 1481–1491 (2022). 10.1038/s41593-022-01176-5

7 Lopes-dos-Santos, V., Conde-Ocazionez, S., Nicolelis, M. A., Ribeiro, S. T. & Tort, A. B. Neuronal assembly detection and cell membership specification by principal component analysis. PLoS One 6, e20996 (2011). 10.1371/journal.pone.0020996

8 Lopes-dos-Santos, V., Ribeiro, S. & Tort, A. B. Detecting cell assemblies in large neuronal populations. J Neurosci Methods 220, 149–166 (2013). 10.1016/j.jneumeth.2013.04.010

9 Kruskal, P. B., Stanis, J. J., McNaughton, B. L. & Thomas, P. J. A binless correlation measure reduces the variability of memory reactivation estimates. Stat Med 26, 3997–4008 (2007). 10.1002/sim.2946

