## Supplemental Material for "Hierarchical encoding of reward, effort and choice across the cortex and basal ganglia during cost-benefit decision making"

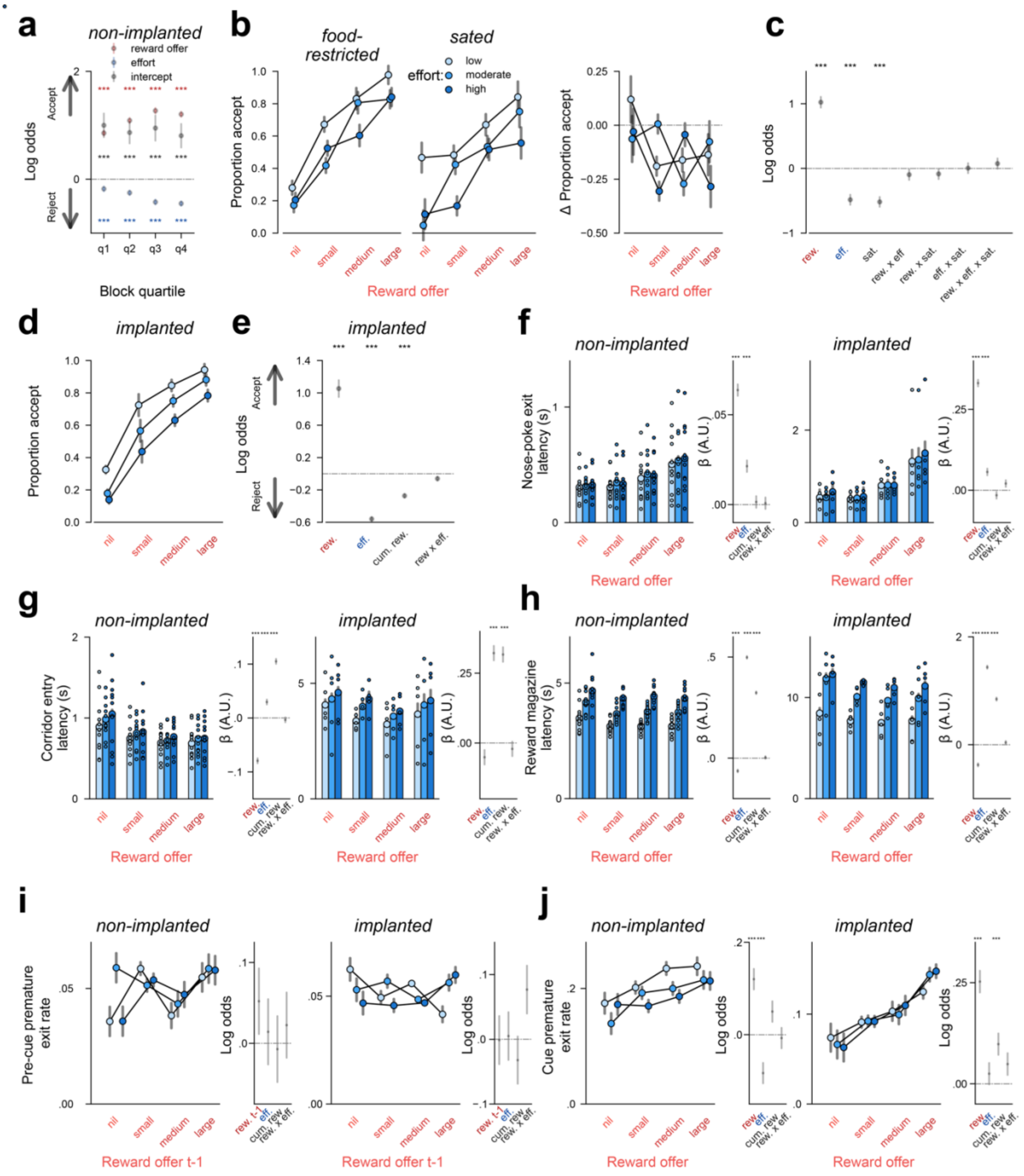


**Supplemental Figure. 1 Choice on a rat accept/reject choice paradigm is shaped by the costs and benefits associated with the offer. a,** weightings of reward and effort within quartiles of each effort block. Data are depicted as regression coefficients for reward (red) and effort (blue) ± SEM. ***, p < .01, corrected for family-wise error rate. **b,** Psychometric curves depicting rates of accepting offers as a function of reward (x-axis) and effort (black lines dotted with coloured scatter) from our within-subject outcome devaluation experiment (n=6). Data are depicted separately for rats' food-restricted and sated state on the left and right, respectively, with the difference (delta) scores between the sated and food-restricted proportions depicted on the right. **c,** weightings of experimental variables on behavioural choice from a binomial mixed effects' model fitted on data from the outcome devaluation experiment (n=6). **d,** Implanted rats' (n=4, sub-selected from the initial cohort of 12) rates of accepting offers as a function of reward and effort. **e,** weightings of task variables on implanted rats' choice from a binomial mixed effects' model. **f-h,** mean nose-poke exit (f), corridor entry (g), and reward magazine (h) latencies (left panels) and associated coefficient weightings from general linear mixed effects' models (right panels, online methods). Nose-poke exit latencies were defined as the interval between cue onset and nose-poke exit, while corridor and reward magazine entry latencies were calculated as the time duration between nose-poke exit and closest corridor light-gate or reward magazine entry, respectively. Bars and larger scatters represent mean ± within-subject SEM, while small scatter dots represent per-subject median values. Colour-code is expressed as per panel **b**. **i-j,** mean pre-cue and cue premature nose-poke exit rate, respectively, and associated coefficient weightings from a binomial hierarchical model (online methods). Generalised linear mixed effects' models in i and j were fitted without nil reward trials. ***, $p < .01$, corrected for family-wise error rate using the Benjamini-Hochberg (BH) method.

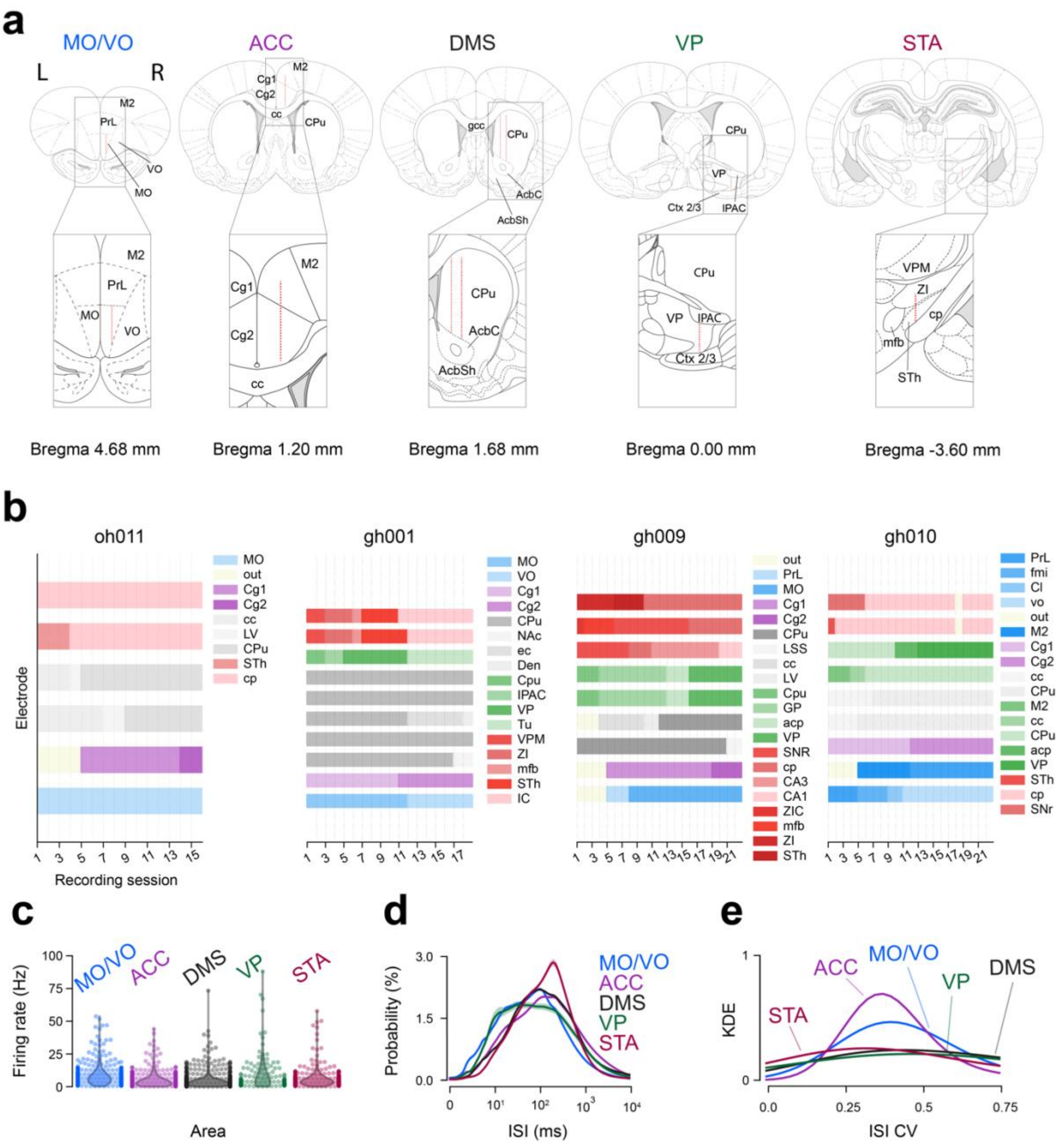


**Supplemental Figure 2. Anatomic localisation and basic firing properties of single units recorded across five frontal-basal ganglia regions. a,** planned stereotrode tracts. Stereotrode trajectories are depicted as dashed red lines superimposed on adapted atlas schematics on whole coronal sections (above) and zoomed insets (below; adapted from Paxinos and Watson 2006). **b,** Estimated targeting data are displayed from each identified electrode tract (bars) within each animal (panels) across all acquired recording sessions (x-axis). Electrode tract targets are colour-coded by the targeted area (blue - MO/VO; purple - ACC; grey - DMS; green - VP; red - STA). **c,** average firing rates of each identified neuron (dots) and kernel density estimates (KDEs) of the neuronal population (violin bodies) coloured by the targeted area. **d,** histogram of average inter-spike interval (ISI) values of neurons in each region. Values are depicted as the population mean probabilities (dark lines) ± SEM (shaded areas). Histograms values are smoothed with a gaussian kernel with a standard deviation of 1.0. **e,** KDEs of neuronal coefficients of ISI variation (lines, colour-coded by area). AcbC - nucleus accumbens core; AcbSh - nucleus accumbens shell; cp - cerebral peduncle; CA1/3 - field CA1/3 of the hippocampus; Cg1/Cg2 - cingulate gyrus 1/2, cc - corpus callosum; CPu - caudate/putamen; cp - cerebral peduncle; ec - external capsule; Den - dorsal endopiriform nucleus; fmi - forceps minor of the corpus callosum; M2 - secondary motor cortex; IPAC -

internal nucleus of the posterior limb of the anterior commissure; LSS - lateral stripe of the striatum; VPM - ventral posteromedial thalamic nucleus; ZI - zona incerta; mfb - median forebrain bundle; out - outside the brain; PrL - prelimbic cortex; STh - subthalamic nucleus; Tu - olfactory tubercle.

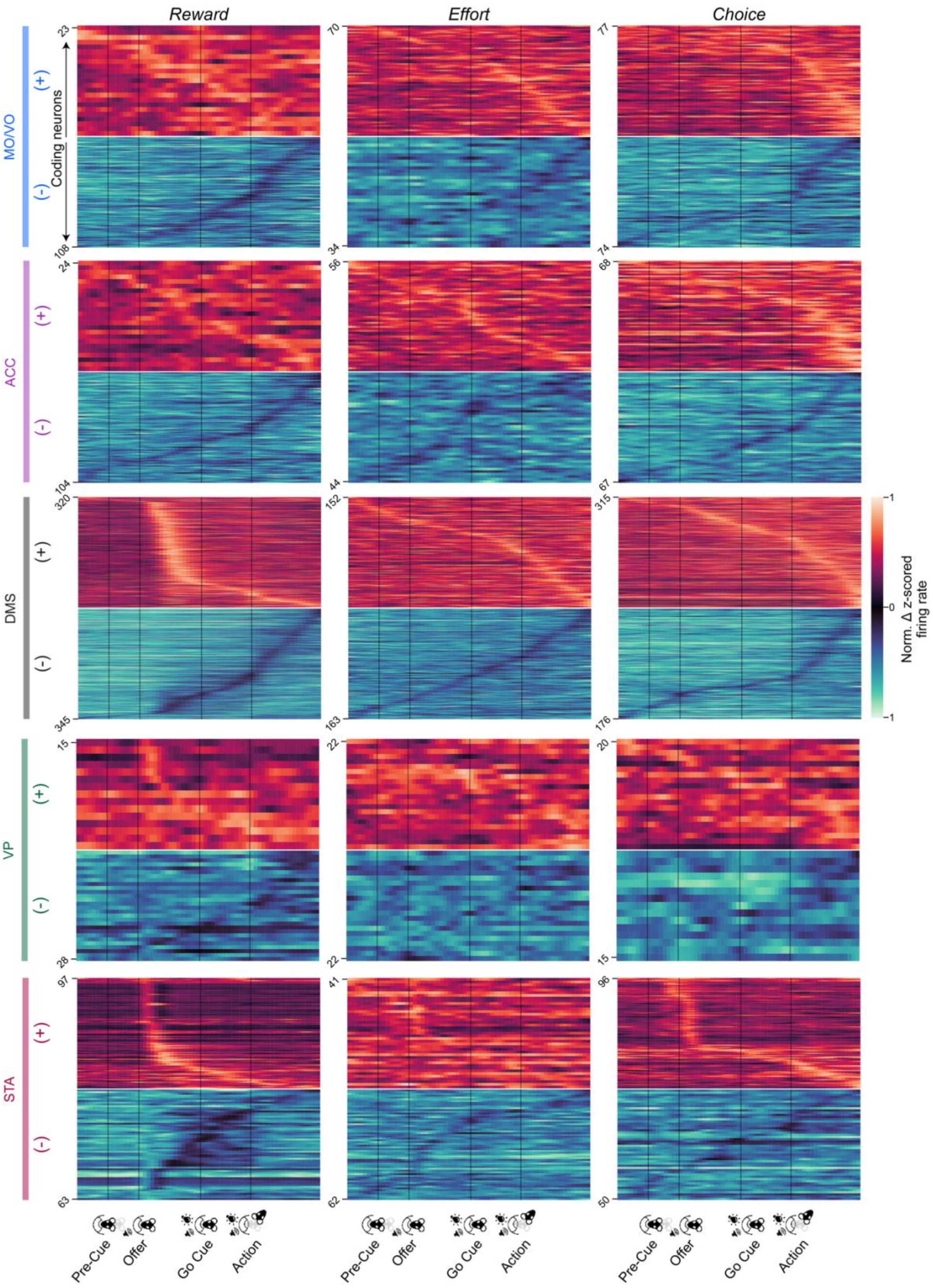


**Supplemental Figure 3. Dynamic single neuronal representations of reward, effort and decision across the frontal-basal ganglia network.** Data are depicted as normalized z-scored firing rate differences for each neuron (row) between the highest and lowest category of each decision variable and sorted by the time bin of peak difference. Heatmap and PSTH data are grouped by positive (above) or negative (below) tuning valence. Neurons were selected on the basis of significant regression coefficients compared to trial-shuffled null distributions (online methods). Time series for each neuron are smoothed with a Gaussian kernel of 1.5 standard deviation.

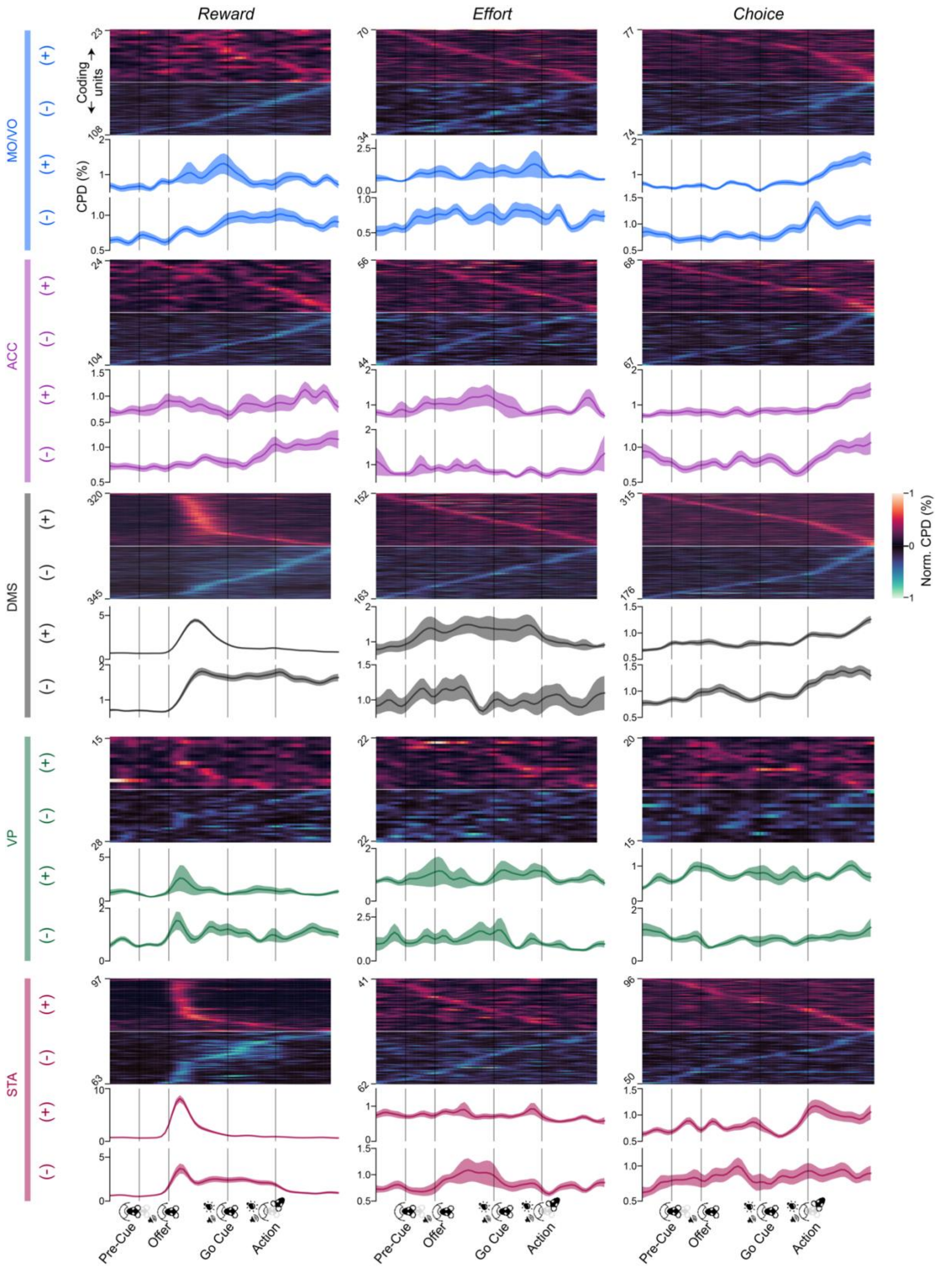


**Supplemental Figure 4. Dynamic and variable single neuronal representations of reward, effort and decision across the frontal-basal ganglia network.** Heatmap data are depicted as CPD values for each significantly coding neuron (row) and sorted by the time bin of peak CPD, while PSTH data represent mean ± SEM across all significantly coding neurons. Heatmap and PSTH data are grouped by positive (above) or negative (below) tuning valence. Neurons were selected on the basis of significant regression coefficients compared to trial-shuffled null distributions (online methods). Time series for each neuron on the heatmaps and for population-average traces are smoothed with a Gaussian kernel of 1.5 standard deviation.

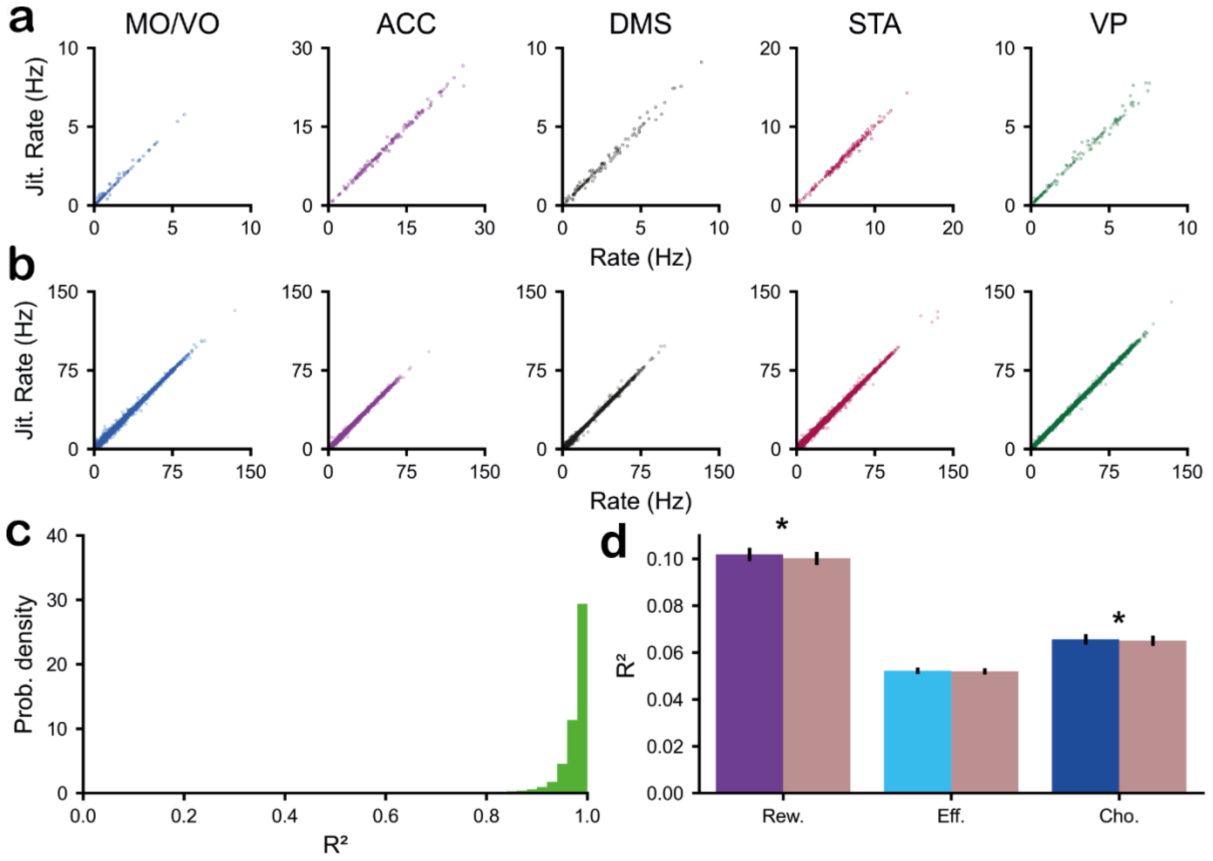


**Supplemental Figure 5. Cofiring assemblies across the frontal-basal ganglia network represent combinations of reward, effort and decision. a**, The trial-by-trial jittered firing rate plotted against the real firing rate for example neurons from the MO, ACC, DMS, STA and VP. **b**, The trial-by-trial jittered firing rate plotted against the real firing rate pooled across all neurons in the MO, ACC, DMS, STA and VP (brain structures ordered as in **a**). **c**, The distribution of the $R^2$ for the correlation between the trial-by-trial jittered firing rate and the real firing rate for each of the recorded neurons. **d**, The $R^2$ between the firing rate of unjittered (brown) as compared to jittered spike trains (blue) in the decision window for reward (p=4.3 x 10-5, Wilcoxon signed-rank test), effort (p = .068, Wilcoxon signed-rank test) and decision (p = .038, Wilcoxon signed-rank test) for significantly coding neurons (significance determined using p-values from linear regressions with firing rate as a dependent variable and reward, effort and choice as independent variables).
